# Dense time-course gene expression profiling of the *Drosophila melanogaster* innate immune response

**DOI:** 10.1101/2020.06.25.172452

**Authors:** Florencia Schlamp, Sofie Y. N. Delbare, Angela M. Early, Martin T. Wells, Sumanta Basu, Andrew G. Clark

**Affiliations:** Molecular Biology and Genetics, Cornell University, Ithaca NY, United States; Statistics and Data Science, Cornell University, Ithaca NY, United States

**Keywords:** *Drosophila melanogaster*, immune response, time course RNA-seq, Granger causality

## Abstract

**Background:** Immune responses need to be initiated rapidly, and maintained as needed, to prevent establishment and growth of infections. At the same time, resources need to be balanced with other physiological processes. On the level of transcription, studies have shown that this balancing act is reflected in tight control of the initiation kinetics and shutdown dynamics of specific immune genes.

**Results:** To investigate genome-wide expression dynamics and trade-offs after infection at a high temporal resolution, we performed an RNA-seq time course on *D. melanogaster* with 20 time points post Imd stimulation. A combination of methods, including spline fitting, cluster analysis, and Granger causality inference, allowed detailed dissection of expression profiles, lead-lag interactions, and functional annotation of genes through guilt-by-association. We identified Imd-responsive genes and co-expressed, less well characterized genes, with an immediate-early response and sustained up-regulation up to five days after stimulation. In contrast, stress response and Toll-responsive genes, among which were Bomanins, demonstrated early and transient responses. We further observed a strong trade-off with metabolic genes, which strikingly recovered to pre-infection levels before the immune response was fully resolved.

**Conclusions:** This high-dimensional dataset enabled the comprehensive study of immune response dynamics through the parallel application of multiple temporal data analysis methods. The well annotated data set should also serve as a useful resource for further investigation of the *D. melanogaster* innate immune response, and for the development of methods for analysis of a post-stress transcriptional response time-series at whole-genome scale.

## BACKGROUND

Upon microbial infection, *Drosophila* launch rapid and efficient immune responses that are crucial to survival. However, immune responses are energetically costly (1) because they draw resources from other physiological processes (2, 3) such as metabolism, reproduction, and environmental stress responses. An excessive or overly prolonged immune response can lead to metabolic dysregulation, causing wasting in mammals and flies (4). Furthermore, it has been shown that allocating resources to the immune system reduces resources for mating (5, 6), and the opposite is also true, where mating reduces survivorship after infection and decreases resistance to infection (7–9). This represents a trade-off where limited resources need to be allocated to either the immune response or reproduction (10). Therefore, we expect that natural selection will act to tune the immune response to strike a balance between the advantage of a rapid and robust ability to fight infection, and the costly side-effects of an over-prolonged immune response. This tuning is likely to be mediated through a series of regulatory and feedback properties of the immune system of the fly.

While gene expression has been examined at several time points after infection in *Drosophila* (11–13), the dynamics of this immune response have not yet been studied with high temporal resolution. A high-resolution time-course analysis can help profile with more certainty the types of expression dynamics that different genes and pathways undergo after infection. Dense and extended time-course sampling of gene expression of the immune response can allow us to distinguish between transient and sustained expression patterns, where expression of genes with a transient response to perturbation will return back to normal after a certain period of time, while expression of genes with a sustained response will remain at a different level of expression compared to pre-perturbation levels. This kind of temporal profiling of the immune response, coupled with computational modeling of gene interaction networks, can also suggest candidates to examine for possible interactions and trade-offs between the immune response and other physiological processes.

Statistical analysis of such high-dimensional longitudinal time-course omics data is not straightforward. While the problems of detecting differentially expressed (DE) genes and inferring gene interaction networks from gene expression data are common in genomics, computational methods have focused primarily on cross-section rather than time-course data. Most popular methods to analyze static RNA-seq data — such as edgeR (14) or DESeq2 (15) — are not ideal for dealing with time-course RNA-seq data since they do not directly model the correlation of genes between successive time points (16, 17). Smooth polynomial or spline based models of temporal dependence in gene expression, such as those employed in Limma-Voom (18) and maSigPro (19, 20), can fail to capture early impulses in stress response situations, as we highlight in this paper. Also, joint network inference of temporal associations among many genes requires tackling high-dimensionality, an aspect that has not received much attention in the literature. Because there is not one consensus method for the analysis of time-course RNA-seq data, it is important to ensure robustness of findings across different types of computational modeling techniques.

In this study, we performed a dense time-course RNA-seq analysis of the *Drosophila* transcriptional response to commercial *E. coli*-derived crude lipopolysaccharide (LPS), which activates the Imd pathway (21), to better understand the dynamics of activation and resolution of the innate immune response. Flies were sampled over 5 days generating a total of 20 time points post-LPS injection with an additional time point pre-injection as a baseline control. We analyzed the resulting longitudinal RNA-seq dataset using a broad range of statistical methods, including a cross-sectional and a dynamic method for differential expression (DE), clustering, and multivariate Granger causality (22), a method to investigate lead-lag relationships among DE genes. We found that commercial LPS exposure has a major impact on the expression of not only immune genes, but also genes involved in metabolism and replication stress. Clustering analysis showed that both the onset and persistence of expression changes varied across these DE genes. Clustering analysis further suggested a role in the immune response and circadian rhythm for several previously uncharacterized genes. Finally, throughout our analyses we observed a theme of interplay and trade-off between the immune response and metabolism.

## RESULTS

### High-resolution profiling of gene expression after Imd stimulation

To generate a full transcriptional profile of gene expression dynamics in *Drosophila melanogaster* after Imd stimulation, we injected adult male flies with commercial *E. coli*-derived crude lipopolysaccharide (LPS). While pure *E.coli* LPS does not induce an immune response in *Drosophila*, the peptidoglycan contamination in crude LPS preparations consistently activates the Imd pathway (21). This was also confirmed using qPCR (see **Materials and Methods**). In the remaining text, we refer to this treatment as “Imd stimulation”. Using commercial LPS instead of living bacteria gives the advantage of avoiding the confounding effects of a growing and changing population of pathogens, and of the mechanisms the bacteria use to circumvent immune responses (23).

Flies were sampled in duplicate for a total of 21 time points throughout the course of five days, which includes an uninfected un-injected baseline sample as control at time zero, and 20 time points after injection. Since this is a perturbation-response experiment, denser sampling occurred at early time points (16), with the first 13 time points taken within the first 24 hr (1, 2, 3, 4, 5, 6, 8, 10, 12, 14, 16, 20, and 24 hr). Sampling is also essential at later time points to know how long it takes to return to baseline, and to differentiate between transient and sustained responses (16). For this reason, sampling continued until day 5 after Imd stimulation, although more sparsely (30, 36, 42, 48, 72, 96, 120 h) (**Figure 1A**). For this dataset we obtained 41 high-quality libraries with an average of 23.5 million mapped reads per sample. After normalization of libraries, only genes with more than 5 counts in at least 2 samples were kept, leaving 12,657 genes for further analysis.

**Figure 1.**
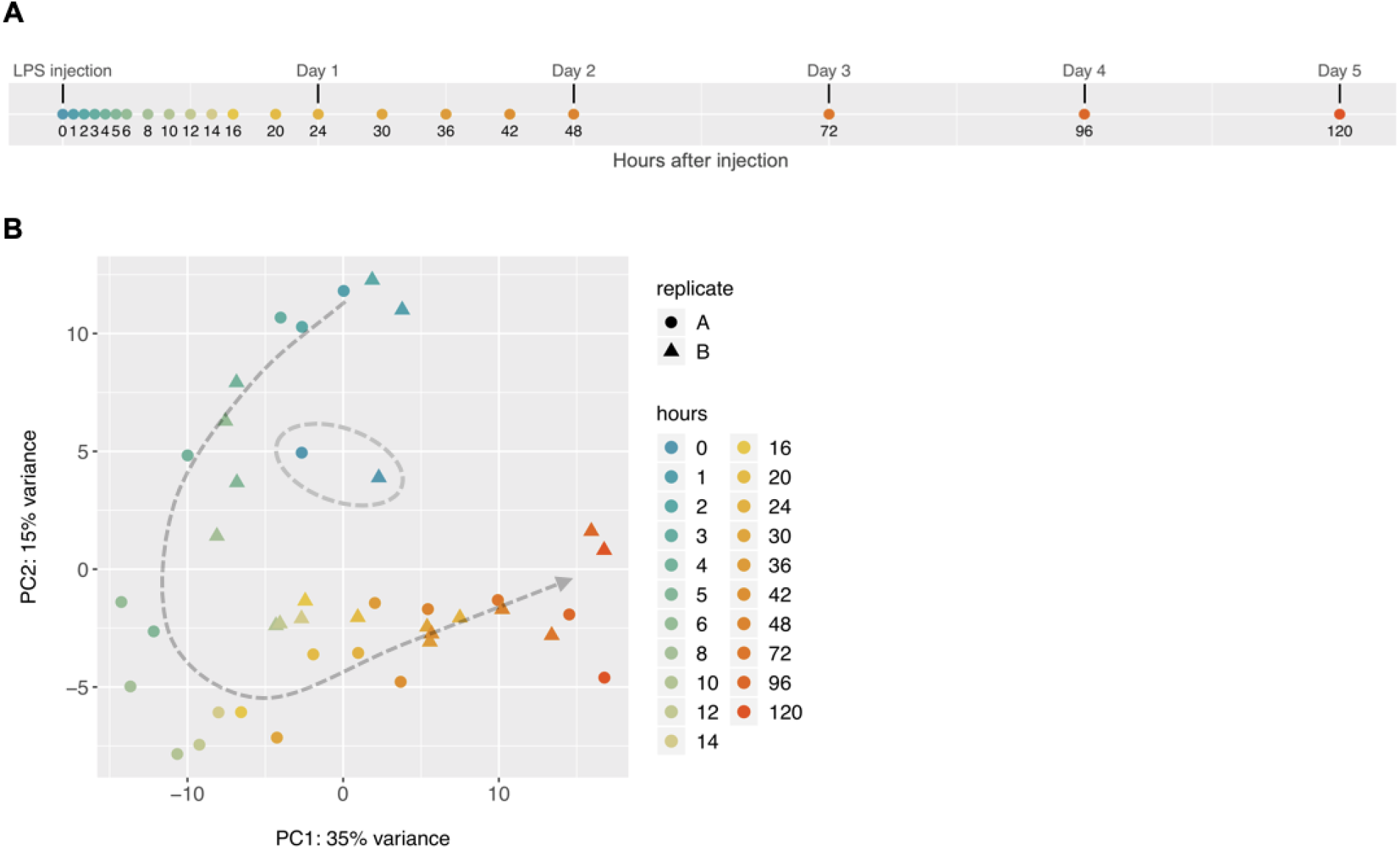
Transcriptional profiling of *Drosophila* immune response. (**A**) Timeline of 21 time points, including un-infected un-injected sample as control at time 0. Sampling was denser in the first 24 hr and continued — although more sparsely — until day 5 (120 h). (**B**) Principal component analysis (PCA) of the top 500 genes with highest row variance across all time points shows a coordinated change of gene expression over five days. Both replicates are shown for all samples except for the time point at 3 h, where one replicate was excluded from the analysis during RNA-seq data processing. The two samples in blue clustering in the middle (marked with grey dashed circle) correspond to the control time point (0 h). All other time points from 1 to 120 hr show a horseshoe temporal pattern around the controls.

Principal components analysis (PCA) on the 500 genes with highest row variance across all time points revealed a horseshoe temporal trend, with the control samples clustering in the middle, and the post-injection time points following a horseshoe-shaped track, consistent with a pattern of many genes displaying a coordinated change over the five-day interval (**Figure 1B**). This type of “horseshoe” or arch temporal trend in PCA has been seen in other time-series experiments (18, 24–26), and is commonly seen in spatial population genetic variation (27) and in ecological gradient data that varies in a non-linear manner (28). PC1, PC2, and PC3 captured 35, 15, and 14.5% of the variance in gene expression respectively, and the first six PCs account for over 80% of the total variance in the data.

Proper normalization of the data was assessed by confirming the behavior of known *Drosophila* housekeeping genes across time (Qiagen Housekeeping Genes RT^2^ Profiler PCR Array and (29)). As expected, housekeeping genes showed little change across time (**Figure S1A**). The success of the Imd stimulation was confirmed by the immediate up-regulation of known immune response genes within the first time points (**Figure S1B**).

### Spline modeling and pairwise comparisons identify 951 genes that are differentially expressed over time following Imd stimulation

To identify genes whose expression levels were significantly altered across the time course, we employed two methods. First, we used gene-wise linear models to fit cubic splines with time, on both the first 8 hr and first 48 hr after commercial LPS exposure. Second, because we noticed that certain expression patterns were not adequately described using cubic splines (as discussed below), we also characterized the temporal patterns of expression by estimating the differential expression of every gene at each time point, from 1 to 48 hr, compared to the un-infected un-injected control samples at time zero (also referred to as baseline).

Cubic spline fits identified a total of 411 DE genes, based on a 5% False Discovery Rate (FDR) using the Benjamini-Hochberg method (30) (**Table S1**). Of these 411 genes, 31 genes were detected only using short spline fits on the first 8 hr post-injection. Long spline fits on the first 48 hr post-injection identified 363 genes, and 17 genes were identified using both short and long spline fits (**Figure 2A**). Long spline fits excelled at identifying gradual changes and global patterns, such as the ones shown by genes *Gale* and *Galk* (**Figure 2B**; note that these genes are mentioned here only to illustrate the spline fit to expression dynamics, and not for their biology). However, long spline fits failed to detect early impulse patterns, such as those observed in the known immune response genes *AttA* and *DptB* (**Figure 2C**), which were better captured by short spline fits on the first 8 hr post-injection. Still, even short spline fits failed to identify additional known immune genes with early impulse patterns, such as *CecC and CecB*. We also fit third degree polynomials using the R package maSigPro (19, 20). This approach identified many DE genes that had been selected using spline modeling (**Figure S2**), but similarly failed to adequately describe early impulse patterns. Based on these observations, we also used pairwise comparisons to identify additional DE genes whose trajectories were not well described using cubic splines. Pairwise comparisons identified 729 DE genes that were significantly (FDR < 0.05) up- or down-regulated by a log_2_ fold change (log_2_FC) of at least 1 (i.e.: 2 times higher than baseline) in at least one time point throughout the first 48 hr after injections (**Table S1**). Within this gene set, there were 214 genes that were up- or down-regulated by at least 2 log_2_FC (4 times higher than baseline), in at least one time interval after injection (**Figure S3**, **Table S1**). Of these 214 genes, 91 “core” DE genes underwent at least a 2 log_2_FC in expression in at least two time intervals after injection, with a more stringent FDR < 0.01 (**Figure 3A**). Among the most strongly induced genes were known immune genes *DptB, AttC, Mtk, Dro, DptA* and *edin* (bottom of **Figure 3A** and **Figure S4A**). These genes underwent an expression change of approximately 5 log_2_FC (32 times higher than baseline) and remained elevated up until 48 hr after Imd stimulation. Further investigation of the 91 core genes showed that the number of up-regulated genes was much higher than the number of down-regulated genes across all time points (**Figure 3B**). Eleven of the up-regulated genes at each time point were known immune genes, as identified by a list of immune genes curated in Early *et al.* (31). Within these 91 core DE genes, we also found circadian rhythm genes *period* (*per*), *timeless* (*tim*), *takeout* (*to*), and *vrille* (*vri*), which when plotted against time exhibit the classic 24 hr periodic expression of the circadian rhythm (**Figure S4B**).

**Figure 2.**
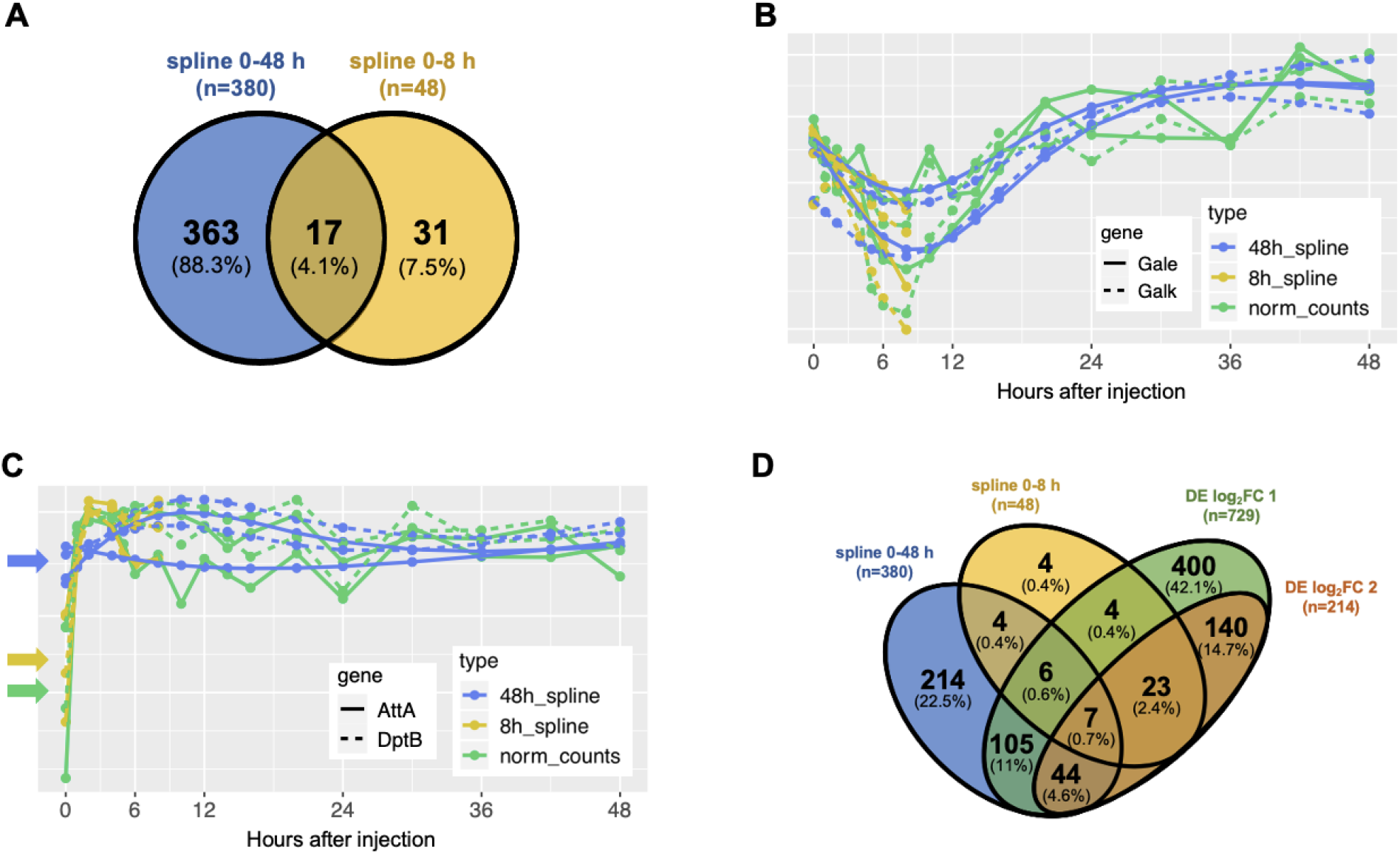
Identification of time-dependent genes. (**A**) Genes that significantly change in expression across time according to spline analysis in the first 8 hr (yellow) vs 48 hr (blue). (**B**) Spline modeling of genes *Galk* and *Gale* when using first 48 hr (blue) and first 8 hr (yellow) compared to the pattern of normalized counts (green). Spline modeling over 8 hr misses the main change in pattern. (**C**) Spline modeling of two immune genes (*AttA* and *DptB*) when using first 48 hr (blue) and first 8 hr (yellow) compared to the pattern of normalized counts (green). Spline modeling over 48 hr smooths out the early impulse signal, drastically changing the inferred expression at time zero (indicated with arrows). (**D**) Comparing results from spline analysis (over 48 hr in blue and over 8 hr in yellow) vs. results from differential expression analysis (|log_2_FC| > 1 in green and |log_2_FC| > 2 in orange) at FDR < 0.05.

**Figure 3.**
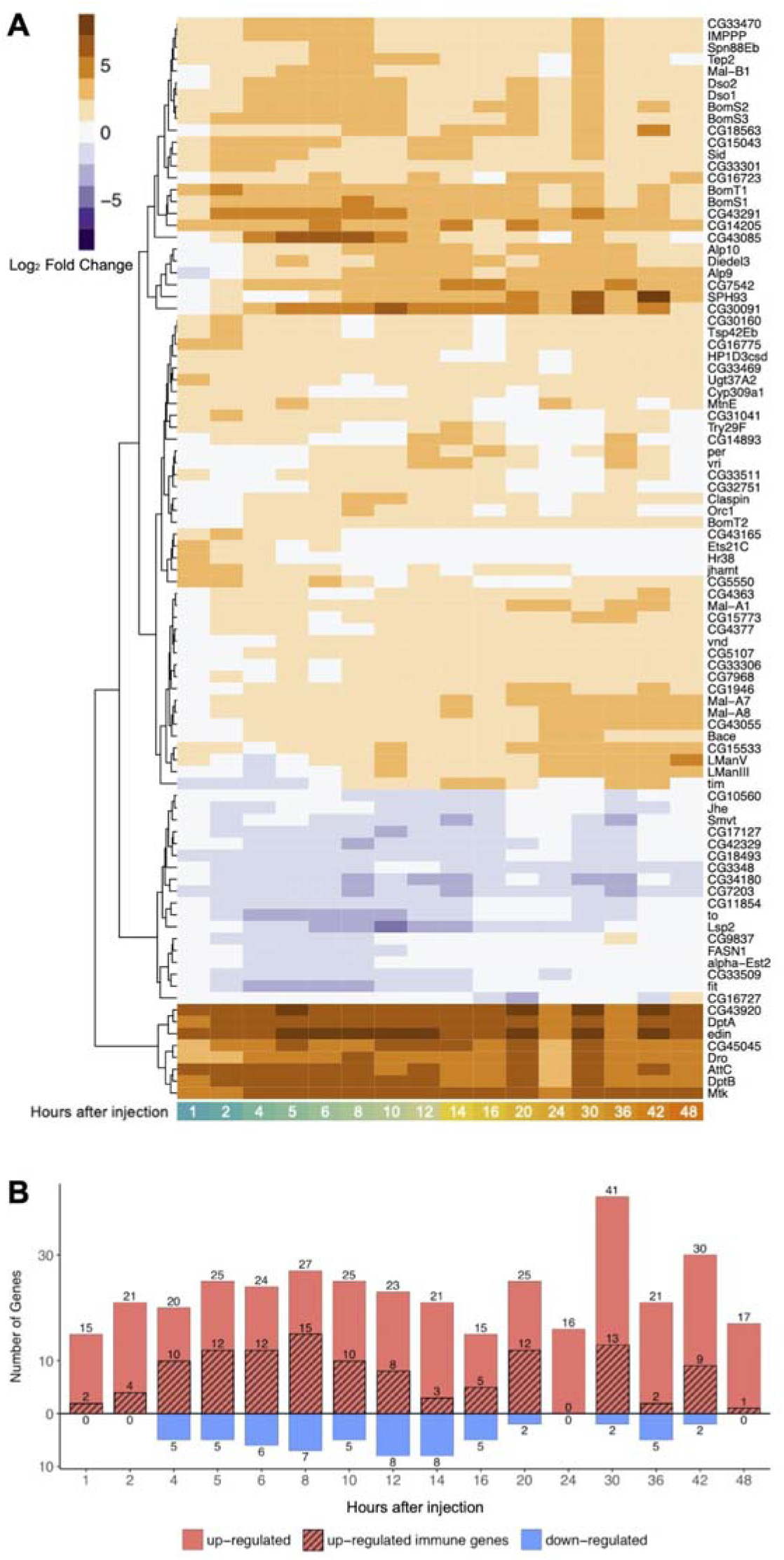
Dynamics and functions of genes with changing expression patterns over time. (**A**) Heatmap of gene expression changes. Up-regulated genes are shown in orange, down-regulated genes are shown in purple. These 91 genes were selected based on FDR correction of 0.01 and a |log_2_FC| > 2 in at least two time points across 48 hr. The genes were ordered using Euclidean distance. (**B**) Number of significantly up- and down-regulated genes, from the core 91 DE genes, at each timepoint (in red and blue, correspondingly). Known immune genes are shaded over red. No down-regulated immune genes were observed.

Of a total of 951 DE genes, 189 genes were identified as differentially expressed using both pairwise methods and spline modeling, but 762 out of 951 genes were identified using only one of these methods, indicating the importance of using complementary methods for the analysis of time course RNA-seq data (**Figure 2D**).

### Gene Ontology and Gene Set Analysis demonstrate a divergence in expression between immune and metabolic processes after Imd stimulation

To understand the biological functions of genes whose expression is influenced by Imd stimulation, we performed both a Gene Ontology (GO) and Gene Set Analysis. GO analysis is a useful tool to illustrate the functions of genes with significant differential expression over time, in this case 951 DE genes selected using spline fitting and/or pairwise contrasts. However, focusing only on the top-scoring genes can lead to missing biologically relevant signals from genes with modest expression changes. Furthermore, GO analysis does not take into account expression changes over time. Both of these limitations are addressed by Gene Set Analysis, which searches for enriched pathways (Gene Sets) across all 12,657 genes in the dataset, guided by their log_2_ fold changes for all available time points.

GO analysis of the 951 DE genes using PANTHER identified a significant (FDR < 0.05) overrepresentation of GO terms related to the immune and stress response, carbohydrate, carboxylic acid and lipid metabolism, and proteolysis. Immune response related genes included Attacins (*AttA*, *AttB*, *AttC*), Diptericins (*DptA*, *DptB*), Cecropins (*CecB*, *CecC*), Bomanins (*BomS1, BomS2, BomS3, BomBc1*), genes encoding Daisho peptides (*Dso1, Dso2*), *IMPPP* (also called *IM10*), *Drosocin* (*Dro*), *Drosomycin* and Drosomycin-like genes (*Drs*, *Drsl1*, *Drsl2*, *Drsl3*), *Metchnikowin* (*Mtk*), Peptidoglycan Recognition Proteins (*PGRP-SB1*, *PGRP-SD*), *Diedel*, *Relish* (*Rel*) and *elevated during infection* (*edin*), among others. DE genes known to respond to stress included Turandots (*TotA*, *TotC*, *TotM*) and Heat Shock proteins (*Hsp70Aa*, *Hsp70Ab*, *Hsp70Ba*, *Hsp70Bb*, *Hsp70Bbb*, *Hsp70Bc*).

Of the 951 DE genes, we identified 20 genes that encode known or putative transcription factors, based on the FlyTF database (32) (**Table S2**). Seven of these twenty genes have a fast impulse of up-regulation, reaching their maximum expression in the first two hours following Imd stimulation (*Rel*, *Dif*, *CrebA*, *luna*, *Ets21C*, *Hr38* and *stripe;* **Figure 4A**). *Rel* and *Dif* encode downstream components of the Imd and Toll pathways respectively, both involved in the activation of the immune response (33–36). Of these two transcription factors, *Rel* undergoes the strongest up-regulation, consistent with activation of the Imd pathway by Gram-negative peptidoglycan in commercial LPS, and remains up-regulated at 1 log_2_FC even after 48 hr. *Ets21C* encodes a stress-inducible transcription factor, and *Hr38* and *stripe* are the two most robust activity-regulated genes (ARGs, defined as genes that are rapidly induced upon stimulation of neurons, mostly within an hour) in *Drosophila* (37). Three genes encoding transcription factors had oscillating expression patterns over time and are involved in the regulation of the circadian clock (*vri*, *clk*, *Pdp1;* **Figure S5A**; (38, 39)). One gene of interest, *p53*, involved in the response to genotoxic stress (**Figure S5B**; (40)), reached its maximum up-regulation later, at 6 hr after injection.

**Figure 4.**
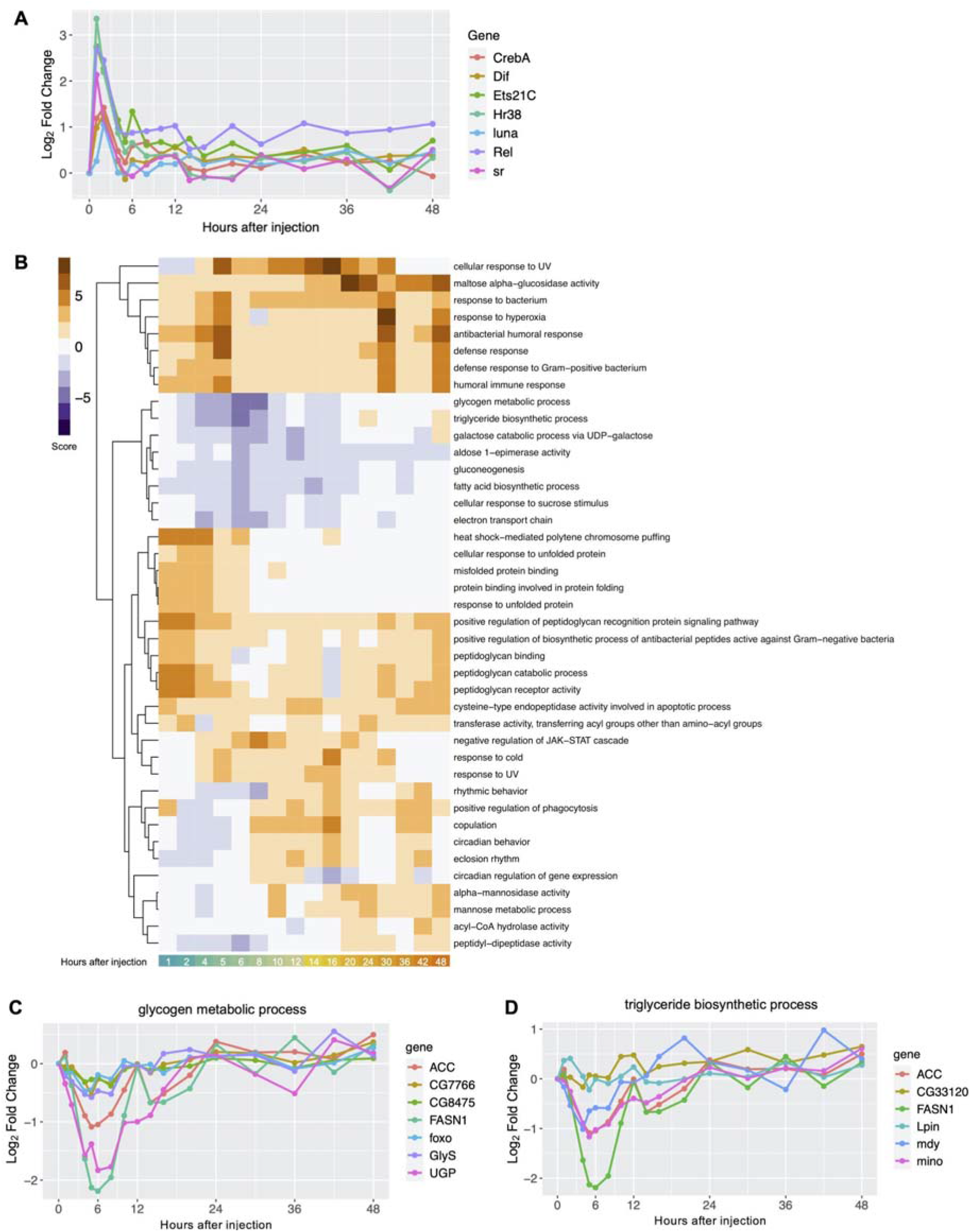
Dynamics and functions of genes with changing expression patterns over time. (**A**) Temporal dynamics of DE transcription factors show that many transcription factors reach their maximal expression 1-2 hr after Imd stimulation. **(B)** Heatmap showing most up- and down-regulated pathways (orange and purple respectively) through the first 48 hr post-injections (absolute pathway score > 2.5 and *P*-value < 0.05 in at least one time point). (**C-D**) **Gene Set Analysis identifies up- and down-regulated pathways.** Selected significantly down-regulated metabolic pathways with corresponding gene memberships.

Overall, the GO analysis indicates that the flies manifest a robust immune response, as the gene expression changes are consistent with known expression profiles of immune response deployment in *Drosophila* (11, 12). In addition, the GO analysis demonstrates that the response to Imd stimulation also affects metabolic homeostasis.

Supporting these results, Gene Set analysis across all 12,657 genes and all time points showed that the top up-regulated pathways were all related to immune response, defense response to bacteria, and peptidoglycan functions (**Figure 4B**). Within these we found pathways related to defense response against both Gram-negative and Gram-positive bacteria. While the commercial LPS used for injections is derived from the outer membrane of Gram-negative bacteria, which activates the Imd pathway, the injections themselves also result in septic injury, which is known to activate both Gram-positive and Gram-negative immune pathways (Toll and Imd pathways correspondingly) (41). Among down-regulated pathways we found many metabolism-related functions. Three of these pathways (glycogen metabolic process, triglyceride biosynthetic process, and gluconeogenesis) are highlighted in **Figure 4C-D** and **S5**.

The glycogen pathway down-regulation pattern was driven by genes *Fatty acid synthase 1* (*FASN1*), and *UGP*, which encodes a UTP--glucose-1-phosphate uridylyltransferase (**Figure 4C**). Down-regulation of the triglyceride pathway was driven by *FASN1* and *minotaur* (*mino*), a glycerol-3-phosphate 1-O-acyltransferase (**Figure 4D**). Finally, the gluconeogenesis pathway down-regulation was driven by *fructose-1,6-bisphosphatase* (*fbp*), a rate limiting enzyme for gluconeogenesis (42) (**Figure S6**). These metabolic genes reached their lowest expression within the first 6 hr after Imd stimulation, and mostly recovered to baseline levels by hours 12-24.

### Clustering of temporal profiles highlights differences in the initiation and shutdown of immune and metabolic genes and demonstrates a regular rhythm of circadian clock genes

GO and Gene Set Analysis illuminated functions of genes that respond to Imd stimulation, and indicated a trade-off between immune and metabolic processes. However, both GO and Gene Set Analysis are based on prior knowledge of gene function. Clustering of genes based only on their expression profiles is not influenced by prior annotations. Such an unbiased approach can thus identify responses of poorly annotated genes. In addition, clustering can illustrate how gene expression trajectories differ over time. We performed three analyses to characterize temporal profiles. First, we performed hierarchical clustering based on Pearson correlation on a set of 551 predominant time-dependent genes to identify major expression patterns over time. These 551 genes included the 411 genes identified using spline modeling, and 214 genes with at least a 2 log_2_ fold change in expression as identified using pairwise comparisons (**Table S1**). Second, we performed clustering based on autocorrelation on these 551 genes. As opposed to Pearson correlation or Euclidean distance, an autocorrelation function takes the ordering of time points into account, allowing us to identify more detailed characteristics of gene expression profiles in time series. Third, because circadian rhythm genes were not apparent in the clusters identified using the previous methods, but were expected to be present in our dataset, we used the R package *JTK_Cycle* (43) to identify genes with 24 hr cycling patterns among all genes in the dataset. We were interested in these patterns since the circadian clock is known to regulate the expression of immune genes (44), and in turn, infections are known to influence the flies’ circadian rhythm (45).

First, expression profiles of the 551 predominant time-dependent genes fell into four main hierarchical clusters (**Figure 5A**). Clusters 1 and 2 both had a strong increase in expression after Imd stimulation (**Figure 5B**). Cluster 1 had a more immediate increase in expression, reaching a maximum within the first 2 hr. Cluster 2, on the other hand, reached a maximum expression later, at around 9 h. Cluster 1 showed significant enrichment of GO terms for immune and stress response related processes, and contained Attacins and Cecropins, as well as Heat Shock protein family genes. Cluster 2 was enriched for GO terms for abiotic stimulus response, and contained among others *Bomanins*, *Daisho* genes, as well as genes from the Turandot family (**Figure 5C**). Clusters 3 and 4 were characterized by an initial decrease in expression followed by an increase after 3 hr and 6 hr respectively (**Figure 5B**), with cluster 4 showing a stronger decrease in expression in the early hours after Imd stimulation. These clusters had a significant enrichment of GO terms for biosynthetic, catabolic, and metabolic processes (**Figure 5C**), and their down-regulation again indicates a trade-off between metabolism and the initiation of an immune response.

**Figure 5.**
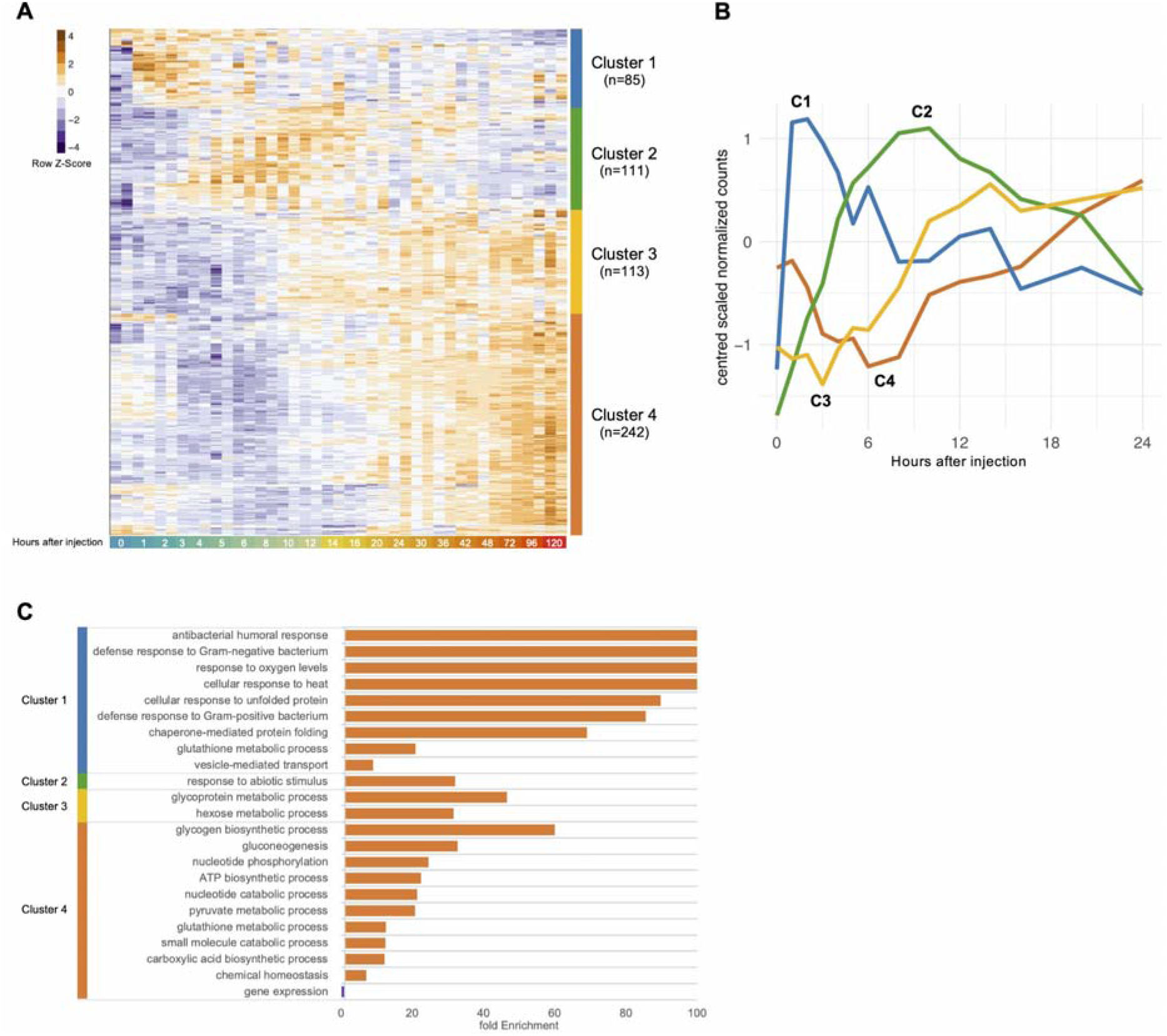
Global dynamics of time-dependent genes show divergent patterns of expression. (**A**) Heatmap of the 551 most predominant time-dependent genes, identified by spline modeling over 48 and 8 hr (FDR < 0.05) and pairwise differential expression (with |log2FC| > 1 and FDR < 0.05). Hierarchical clustering of the genes shows four main clusters characterized by time points in which the genes reach maximum and minimum expression across time. Z-score values of each gene are shown from dark purple (minimum expression across time) to dark orange (maximum expression across time). (**B**) Mean patterns of expression across time for genes within each of the four main clusters during the first 24 hr, displayed by their centered and scaled normalized counts. (**C**) Significant Gene Ontology terms (FDR < 0.05) for over-represented Biological Processes at each cluster.

Our second clustering analysis based on autocorrelation revealed additional differences regarding the initiation and resolution of gene expression after Imd stimulation. First, we identified a cluster of genes with an immediate and sustained up-regulation. This cluster was characterized by a strong early induction with an up-regulation of ∼2.5 to 6 log_2_FC within the first hour (6 to 64 times higher than baseline), reaching a maximum of 6 to 8.5 log_2_FC (64 to 362 times higher than baseline), and maintaining persistent up-regulation of ∼2.5 to 6 log_2_FC throughout 5 days (**Figure 6A**). This cluster contained canonical immune response genes known to be activated downstream of Imd, such as *AttA*, *AttB*, *AttC*, *DptA*, *DptB*, *Dro*, *edin*, *Mtk*, *PGRP-SB1*, *PGRP-SD.* The cluster further contained *IBIN* (*Induced by Infection*), whose exact mode of action is unknown, but whose up-regulation stimulates starch catabolism as part of an immune-induced metabolic switch, likely to make free glucose available to circulating immune cells (46). Finally, this immediate-response cluster also contained *Mtk-like, CG43920* and *CG45045*, which are less characterized transcripts known to be up-regulated after bacterial infection (47).

**Figure 6.**
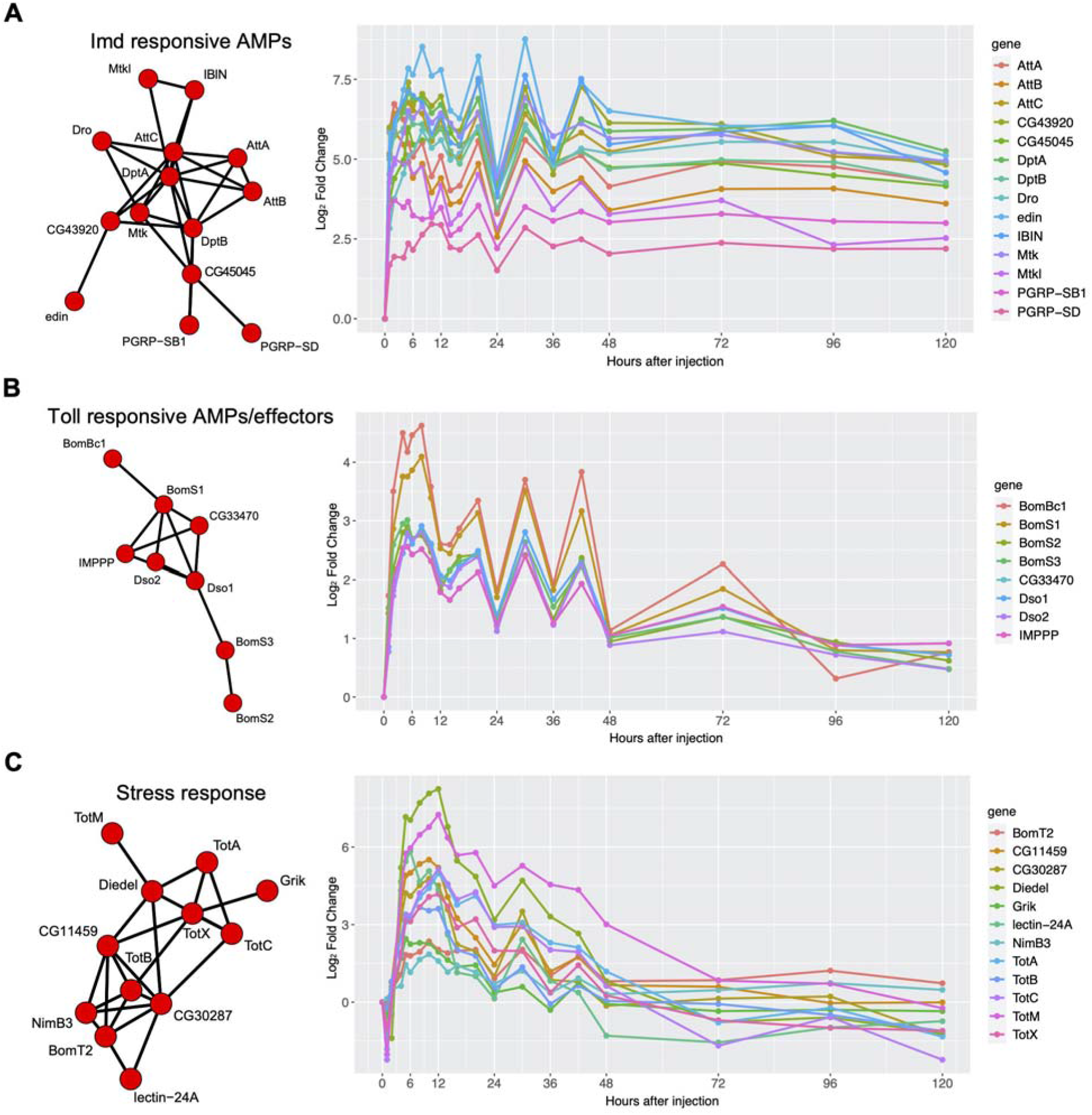
Clusters of genes identified using autocorrelation. Network nodes represent genes; network edges represent the distance between gene autocorrelations, based on ACF analysis using TSclust. (**A**) A cluster containing multiple Imd-responsive genes shows sustained expression after Imd stimulation throughout 5 days (120 h). (**B**) Toll-responsive genes show a transient response to Imd stimulation. (**C**) Other stress response genes return to steady state by day 5 (120 h) post Imd stimulation.

Autocorrelation-based analysis also identified clusters of genes with transient responses to infection (**Figure 6B-C**). One of these clusters was composed of Bomanins (*BomS1, BomS2, BomS3, BomBc1*), *Daisho* genes (*Dso1, Dso2*), *IMPPP* and *CG33470* (**Figure 6B**). The *Bomanins* are located in the 55C4 region of chromosome 2R, confer resistance against certain bacteria and fungi and are regulated by Toll signaling (48). *Dso1 and Dso2* were also reported to be regulated by Toll signaling and aid defense against specific filamentous fungi (49). *CG33470* is an uncharacterized transcript located 3.3 kb downstream of *IMPPP* and might belong to the same open reading frame, as both are sometimes referred to as *IM10* (50), and show nearly identical gene counts in our dataset. This cluster was characterized by an early induction (but not as immediate as the cluster containing known Imd-responsive genes in **Figure 6A**) of ∼2.5 to 3.5 log_2_FC (6 to 11 times higher than baseline) within the first two hours, reaching a max of ∼2.5 to 5 log_2_FC, and returning to a steady state after 3-5 days. Thus, clustering analysis identified effector immune genes partitioned by Imd vs Toll signaling: Imd-regulated genes showed an immediate early sustained up-regulation even after 5 days (**Figure 6A**), while Toll-regulated *Bomanins* and *Daisho* genes had an early up-regulation that eventually returned to steady state levels (**Figure 6B**).

A final cluster illustrated a more complex expression pattern: many genes in this cluster were down-regulated immediately, 1-2 hr after injection, after which they were up-regulated, reaching their maximum expression after 8-12 hr, followed by a return to baseline after 2-3 days (**Figure 6C**). This cluster was composed of genes from the stress-induced Turandot family (*TotA, TotB, TotC, and TotX*) (51) as well as *Diedel*, *Grik*, *lectin-24A*, *NimB3*, *BomT2*, *CG11459*, and *CG30287*. *Diedel* encodes an immunomodulatory cytokine known to down-regulate the Imd pathway. *Grik* encodes a glutamate receptor, and *Lectin-24A* encodes a pattern recognition receptor that mediates pathogen encapsulation by hemocytes (52). *Lectin-24A* has been shown to be down-regulated in the first 2 hr following septic injury and then up-regulated 9 hr after (53), consistent with the pattern we see in our data. *NimB3* is part of the Nimrod gene family, which is involved in phagocytosis (54). *BomT2* is part of the Bomanin gene family (like *BomS1*, *BomS2*, *BomS3* and *BomBc1*, expressed in the previous cluster, **Figure 6B**). *CG11459* encodes a predicted cathepsin-like peptidase induced by bacterial infection and injury (55). *CG30287* encodes a predicted serine protease, a class of proteins that plays roles in immune response proteolytic cascades (56).

Finally, using JTK_cycle, we identified 22 periodic genes with a 24 hr cycle, using a cutoff of Benjamini-Hochberg corrected *Q*-value < 0.05 and amplitude > 0.5 (**Figure 7**). Among them were four well characterized circadian genes, suggesting that their periodicity was not affected by Imd stimulation: *period* (*per*), *takeout* (*to*), *vrille* (*vri*), and *PAR-domain protein 1* (*Pdp1*), as well as eight genes which do not have assigned circadian functions but have evidence of cyclic behavior in previous literature (**Table 1**), and 10 genes, of which 8 are uncharacterized, that have not yet been reported to have cyclic expression outside this study (**Table 1**; *CG10560*, Sgroppino, *CG15253*, *CG15254*, *CG18493*, *CG31321*, *CG33511*, *CG34134*, *CG42329, salt*).

**Figure 7.**
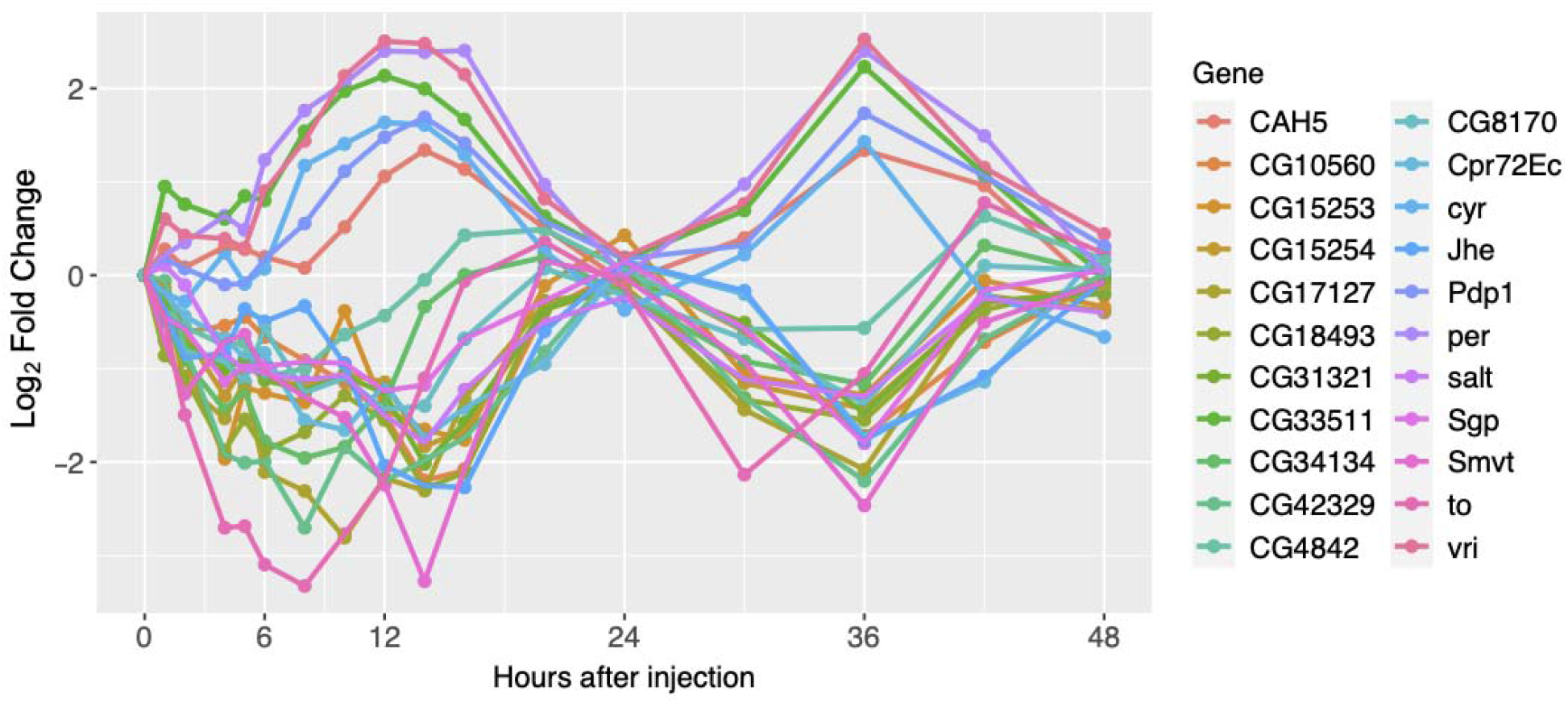
Top 22 genes identified by JTK_Cycle show 24 hr temporal cycling.

**Table 1.**
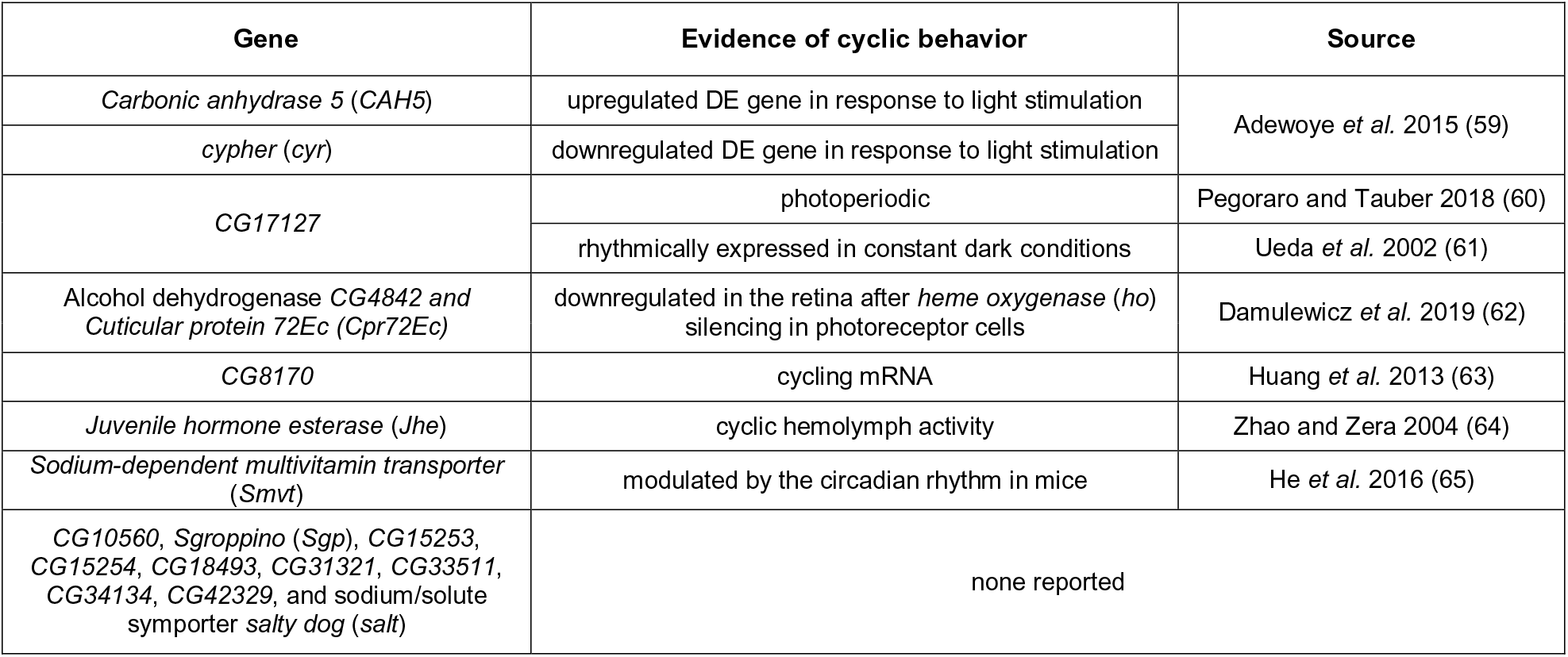
Evidence of cyclic behavior for top genes identified by JTK_Cycle.

Overall, the combination of clustering methods augmented by GO analysis allowed us to identify strong temporal patterns that correspond to early and late induction of immune processes, as well as both transient and sustained responses to infection, which point to a trade-off between the immune response and metabolism. We found that genes that share functions often have similar temporal expression patterns, suggesting co-regulation. This observation further allowed us to assign putative functions to previously uncharacterized genes that cluster together with well-studied genes.

### Gene interaction modeling of lead-lag patterns using Granger causality

Clustering methods based on a single gene’s autocorrelation or cyclicity patterns can detect genes with similar expression profiles. However, these methods are not suitable for seeking causal relationships between genes that manifest in a lead-lag relationship, for instance when high expression of gene A results in a high expression of gene B shortly afterwards. Detecting such lead-lag patterns (**Figure 8A**) is a unique advantage of dense time-course experiments. Granger causality (GC), a statistical method popular in analysis of macroeconomic time series, provides an ideal framework for modeling such patterns and building directed networks among genes. The concept of GC is based on predictability. If the knowledge of the past of one time series improves the prediction of a second one, the first is said to be Granger causal (GC) for the second. *Bivariate* GC analysis between two genes A and B, as described above, does not account for possible confounding effects of other genes C, D, E which can also influence genes A and B (**Figure 8B**). *Multivariate* GC analysis alleviates this problem by explicitly accounting for the effects of the confounding genes by a joint modeling (57, 58), but does not account for high-dimensionality and consequently cannot jointly model hundreds of genes based on tens of data points. We used modern high-dimensional methods (viz LASSO (59) and de-biased LASSO (60, 61)) to address this problem and build lead-lag network models among 258 genes.

**Figure 8.**
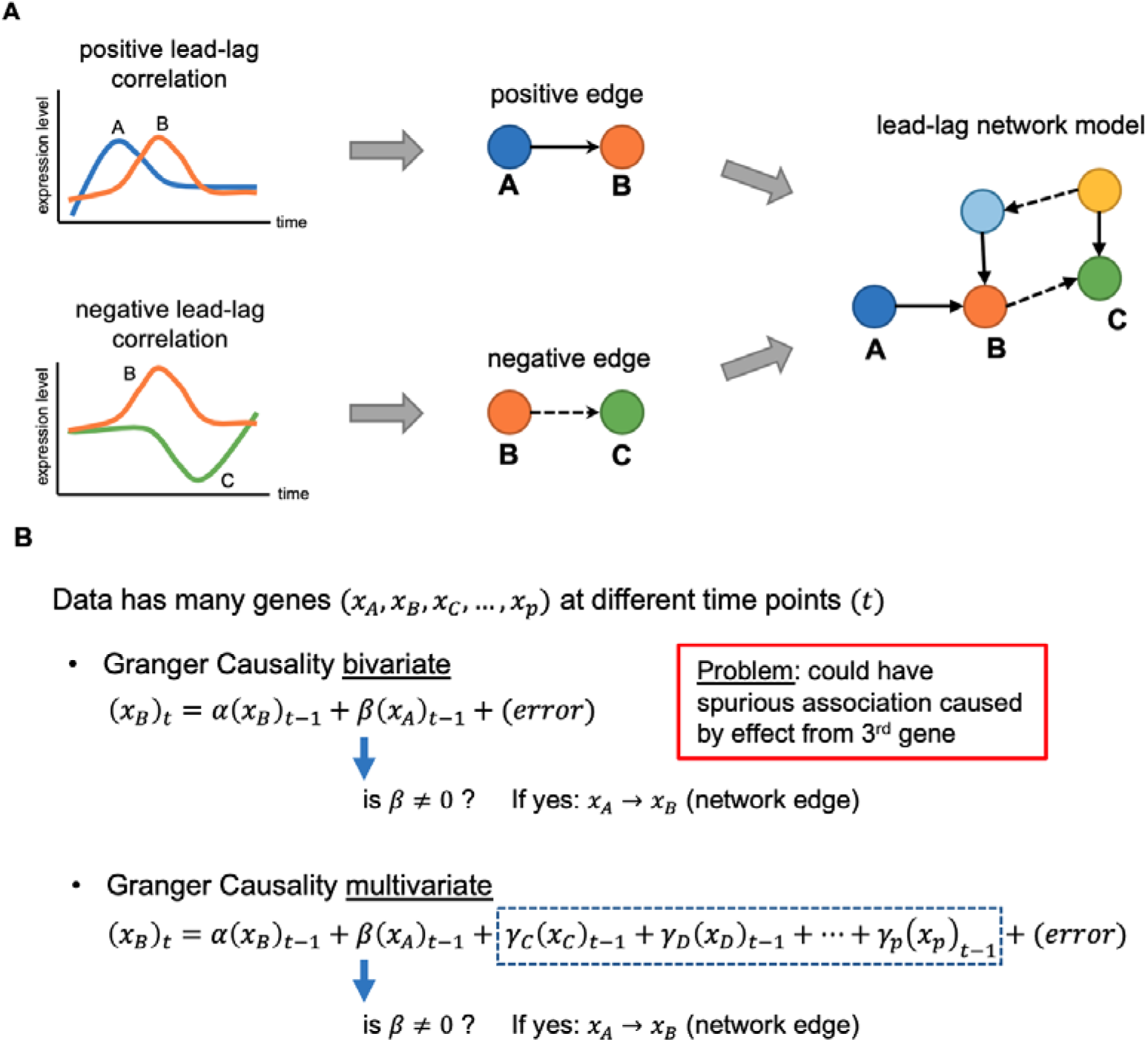
Diagram describing the process of constructing directed networks from Granger causality. **(A)** Lagged correlated expression between two genes (Granger causality) leads to the construction of a directed edge between two genes (nodes), which in turn is used to build directed lead-lag network models of putative interactions among genes. Edges can be positive or negative, based on the sign of lead-lag correlation between the two genes. **(B)** Bivariate associations are calculated between two genes at a time, while multivariate associations adjust for potential indirect association from all other genes in the gene set.

We constructed directed GC edges and networks of putative interactions among a subset of 258 genes (**Table S1**). These genes had a |log_2_FC| > 1 (at least 2 times higher or lower than baseline) across the time course and had available functional annotations. We performed Granger causality analysis on sliding windows of 6 time points on the normalized counts of both replicates using bivariate and multivariate methods (see **Materials and Methods**). We investigated both positive and negative edges, reflecting positive and negative lagged correlations between genes. The overall unfiltered GC network has a multitude of relationships worth exploring, but limitations in the ability to distinguish different types of causality make widespread conclusions from the network challenging. Here, we discuss several examples of subnetworks which illustrate putative functional relationships among genes whose expression changes in response to Imd stimulation.

Based on our interest in identifying trade-offs between biological processes in infected animals, we first constructed a high-quality set of consistently significant GC edges of divergent expression (negative edges). To this end we first filtered the subnetwork by (a) removing all edges with a positive weight, (b) removing all nodes corresponding to cyclic genes identified earlier through the JTK_Cycle method, (c) using only pairs of nodes with significant edges (Benjamini Hochberg FDR < 0.05%) in at least 3 consecutive windows within the first 24 hr of the time course. After filtering, the resulting high-quality GC network contained 51 nodes and 35 edges in 16 connected components (**Figure S7**). This network, by design, should include the most interesting examples of divergent expression changes from our full dataset.

The largest connected component in this network (Component #1) is a multifunctional chain of 6 genes, which connects the down-regulation of four metabolic genes with the up-regulation of two genes that are involved in regulating proliferation and repair (**Figure 9A**). Two of the metabolic genes, *Sorbitol dehydrogenase 1* (*Sodh-1*) and *UGP*, both lead the divergent expression of *Claspin* (both 4 consecutive windows, 2 to 10 and 4 to 12, respectively) (**Figure 9C** & **S8A**). *Claspin* plays a role in DNA replication stress (62). It is known that there is an interplay between host immune systems and replication stress (63). The immune system can detect and respond to replication stress, which is an important feedback loop necessary to remove defective cells (64). Furthermore, the activation of the immune response generates reactive oxygen species (ROS) and reactive nitrogen species (RNS), and can promote chronic inflammation, all of which can trigger DNA damage (65). *UGP* and *fbp* were identified earlier during Gene Set Analysis to drive the down-regulation of metabolic pathways (**Figure 4B** and **4D**), and in this cluster they are both negatively directed by *LpR2* (3 consecutive windows, 6 to 13) (**Figure 9D** and **S8B**). *LpR2* is a lipophorin receptor, known to regulate the innate immune response by clearing serpin protease complexes from the hemolymph through endocytosis (66). Lipophorin is a known humoral factor that contributes to clot formation (67, 68). Finally, *LpR2* is also shown to negatively direct *juvenile hormone acid methyltransferase* (*jhamt*) (4 consecutive windows, 1 to 9) (**Figure 9E**). JHAMT is an enzyme that activates juvenile hormone (JH) precursors at the final step of the JH biosynthesis pathway in insects (69). JH is a known hormonal immunosuppressor in *Drosophila* (70–72).

**Figure 9.**
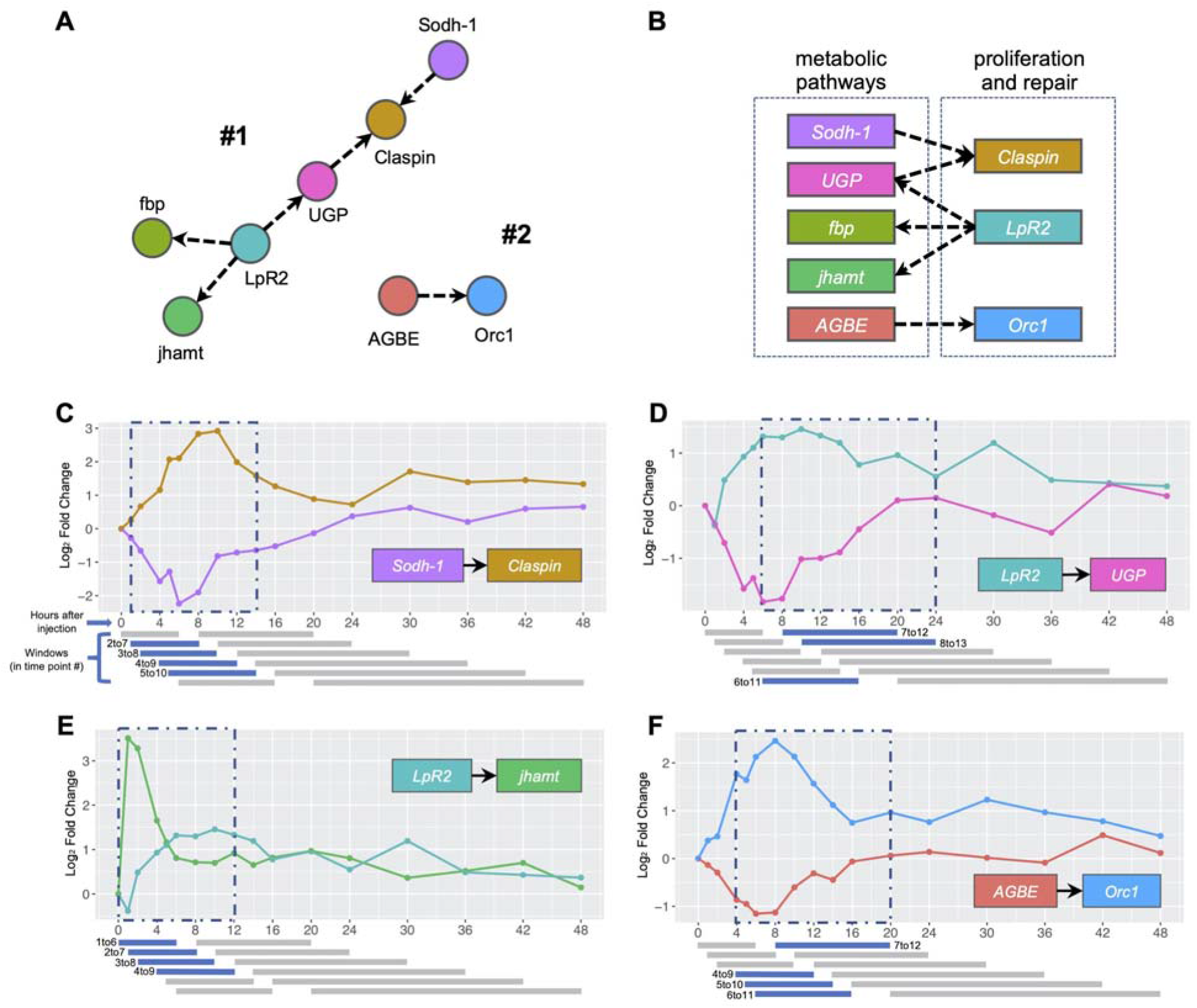
High-quality GC network components and their edges. (**A**) Components #1 and #2 from GC network (**Figure S4**). (**B**) Diagram summarizes interplay between main represented pathways on the selected components. (**C-F**) Selected edges from the components plotted against time. Significant windows colored in blue, non-significant colored in grey. Resulting overall consecutive windows are labeled in blue dashed rectangles. Individual windows represent 6 consecutive time points, but because time points are not at regular intervals, the windows have different time ranges, but identical numbers of samples.

Interestingly, *Claspin* was identified to be part of the same pathway as *Orc1* in our previous Gene Set Analysis, showing similar patterns and window of up-regulation (mitotic DNA replication checkpoint pathway, **Figure S9**). In our network, *Orc1* is part of an isolated edge with metabolic gene *ABGE* (Component #2, 4 consecutive windows, 4 to 12) (**Figure 9A** and **9E**). These prioritized subnetwork components suggest an interplay between metabolic pathways and other pathways such as proliferation and repair (**Figure 9B**), motivating follow-up studies to determine which pathways might be regulating and trading off with each other in the hours following Imd stimulation.

In addition to these purely negative edges, we detected highly significant positive and negative edges among circadian rhythm genes. These included *cryptochrome* and *Smvt* (6 consecutive windows, 6 to 16) (**Figure S10A**), *vrille* and *takeout* (4 consecutive windows, 9 to 17) (**Figure S10B**), *period* and *takeout* (4 consecutive windows, 9 to 17) (**Figure S10C**), and *Smvt* and *takeout* (4 consecutive windows, 9 to 17) (**Figure S10D**). *Smvt* is predicted to encode a sodium-dependent multivitamin transporter, and *takeout* influences feeding behavior (73, 74). Metabolic processes and feeding are known to be under circadian control (75). In addition, So *et al.* (76) reported that *takeout* is regulated by the circadian clock, but with a phase shift relative to *period*. This pattern is clearly visible in our dataset and was correctly identified using Granger causality. This shows that Granger causality can be used to infer gene dependencies/interactions using global gene expression behavior.

Finally, among genes connected only by positive edges, we identified an edge from *period*, a regulator of the circadian clock (77, 78), to *Rhodopsin 5*, which encodes a G-protein-coupled receptor involved in phototransduction (**Figure S11A**). *Rh5* mRNA levels are known to demonstrate a cyclic pattern (79), indicating regulation by the circadian clock. We further identified positive edges between genes that are likely co-regulated. These included edges between up-regulated genes that respond to NF-κB signaling, such as edges from genes encoding peptidoglycan recognition receptors *PGRP-SD* and *PGRP-SB1*, (regulated by Imd signaling), to *DptB* and *AttC* (also regulated by Imd signaling) and to *BomS1*, *Dso1* and *BomBc1* (regulated by Toll signaling) (**Figure S11B-F**). We also observed edges between down-regulated genes, such as from *Hao* (predicted to play a role in lactate oxidation) to *AGBE* (a predicted hydrolase involved in glycogen synthesis) (**Figure S11G**). While the expression of these genes likely responds to similar signals, the observed lags between these genes’ expression profiles suggest that there are differences in their transcriptional control, such as regulation by cofactors, differences in promoter affinity for certain transcription factors, or other variables that influence the rate of mRNA accumulation.

## DISCUSSION

We have produced a dense and high-quality time-course profiling of the *Drosophila* transcriptome response after Imd stimulation through commercial LPS injection, using RNA-seq sampling over 20 time points spanning 5 five days. This profiling provides a high-dimensional dataset, which is available as a resource for the community. We analyzed this dataset using a broad range of statistical methods, including Granger causality, to investigate lead-lag relationships between genes. Because of the high dimensionality, it is not straightforward to analyze as a time series, as illustrated by the partially distinct results of spline fitting and pairwise comparisons. However, using a combination of analytical methods allowed us to identify distinct patterns with high confidence, specifically responses to Imd stimulation with divergent initiation and resolution dynamics, as well as cyclic patterns of gene expression, and patterns of co-regulation and trade-offs. Below, we describe and discuss the main insights from these analyses, as well as limitations and future steps.

### Immune and stress response genes vary in their initiation and resolution dynamics after Imd stimulation

Clusters of genes demonstrated distinct activation kinetics after stimulation of the immune response. This phenomenon has been observed both in fly (11) and mammalian cells (80), but as a result of the dense sampling, our dataset provides a highly detailed view of these initiation dynamics. In addition, because we sampled up to five days post-injection, we could also observe long-term responses to Imd stimulation.

First, a cluster of 13 genes showed the fastest up-regulation within the first 1-2 hr and remained up-regulated during the entire five-day time course (**Figure 6A**). This cluster contained 10 genes known to be regulated by Imd signaling (3 *Attacins*, 2 *Diptericins*, *Mtk*, *Dro*, *IBIN*, *PGRP-SB1* and *PGRP-SD*), as well as *CG43920*, *CG45045 and Mtk-like.* Imd-regulation of these 3 genes has not been experimentally validated, but their co-clustering pattern suggests that they are regulated by the Imd pathway.

Second, a cluster containing Toll-regulated *Bomanins* and *Daisho* genes (81) reached its highest point of expression at 5-8 hr and recovered to a baseline state after 2 to 5 days (**Figure 6B**). The later initiation of Toll-responsive genes relative to Imd-responsive genes is consistent with Tanji *et al.* (82), who reported earlier peak expression of an *Attacin* and *Diptericin* (Imd-responsive) versus *Drosomycin* (mostly Toll-responsive). The up-regulation of Toll-responsive genes might be a response to the wounding that occurred during LPS injection, as Irving *et al.* (83) reported that septic injury with Gram-negative *E. coli* induced responses to both Gram-negative and Gram-positive bacteria, and Boutros *et al.* (11) reported that clean injury experiments induced a set of genes that overlapped with those induced by a septic injury experiment, but with a lower response magnitude, consistent with our observations (**Figure 6A-B**; Imd-responsive genes reach a maximum of 6-8 log_2_FC while Toll-responsive genes reach a maximum of 2.5-4.5 log_2_FC).

Third, stress response genes, among which were members of the Turandot family, reached their highest point of expression at 10-12 hr, following a pattern of delayed response in line with observations by Ekengren *et al.* (51). Similar to the Toll-regulated gene cluster, stress response genes returned to a baseline state after 2 to 5 days (**Figure 6C**).

It is striking that the expression of Imd-regulated genes did not recover to pre-injection levels even after five days. Since we stimulated the Imd pathway by injecting Gram-negative peptidoglycan within commercial LPS, it is possible that the prolonged up-regulation of AMPs and other Imd-regulated genes was due to remaining peptidoglycan in the flies. Alternatively, it might be typical for these genes to be expressed at a higher level for a certain time even after the peptidoglycan has been cleared. Troha *et al.* (47) also observed prolonged AMP up-regulation after bacterial infection, even when levels of bacteria were below detection threshold. On the other hand, Duneau *et al*. (84) reported that seven days after an initial infection with *E. coli*, none of the surviving flies were completely free of bacteria - suggesting suppression rather than clearance of the infection. These observations raise the question of whether one should expect to see a return to the baseline gene expression levels. Rather than returning to a pre-infection state, the prolonged up-regulation of Imd-responsive genes and the transcription factor *Rel* (**Figure 4A**) might be required to prevent damage from dormant bacteria.

### Initiation of the immune response coincides with a down-regulation of metabolic processes

Our dataset showed distinct global dynamics pointing to a divergence in expression between immune and metabolic processes (both carbohydrate and lipid metabolism), with the most up-regulated pathways related to immune and stress responses, and the most down-regulated pathways related to metabolic functions (**Figure 5**).

Metabolic genes reached their maximum down-regulation at 5-8 hr (**Figure 4B-D, 5**). *FASN1*, which showed the strongest down-regulation in both glycogen metabolic process and triglyceride biosynthetic process (**Figure 4C-D**), is a lipogenic gene whose down-regulation might indicate a need to have easily accessible nutrients instead of storing them. Indeed, infections in mammals are known to induce adipose tissue lipolysis (85) and bacterial peptidoglycan is a ligand that stimulates lipolysis as well (86). The gene with the strongest down-regulation in the gluconeogenesis pathway was *fbp* (**Figure S6**), which codes for fructose-1,6-bisphosphatase, the rate limiting enzyme for gluconeogenesis. This gene was significantly down-regulated in a study that reported that *Listeria monocytogenes* infection in *Drosophila* causes a decrease in energy stores, with reduced levels of triglycerides and glycogen (87). Krejcova *et al.* (88) reported that an infection-induced switch to aerobic glycolysis within macrophages coincides with a systemic depletion of glycogen stores and increased blood sugar levels. The divergent dynamics detected in our dataset are thus in agreement with known individual mechanisms characterized in the immune response.

We also observed that expression of metabolic genes and pathways recovered quickly to a baseline state around 12-24 hr after Imd stimulation (**Figure 4B-D, 5**). The speedy recovery of metabolic genes, despite sustained expression of Imd-regulated immune response genes, suggests that the early stages of infection likely involve the greatest trade-offs, at least in response to the commercial LPS injections. These dynamics might differ in flies infected with live bacteria and might also differ depending on the strain of bacteria (47).

We further saw implications of functional interplays using the Granger Causal (GC) network analysis. Main subnetwork components showed significant GC directional edges between down-regulated metabolic genes (such as *Sodh-1*, *UGP*, *fbp*, and *AGBE*) and up-regulated genes with cell proliferation and repair functions (*Claspin*, *LpR2*, and *Orc1*) (**Figure 9**). These results further suggest an underlying interplay between metabolic pathways and proliferation and repair mechanisms such as regulation of DNA replication stress, endocytosis, and clot formation. While functional genetics studies have demonstrated such trade-offs previously (3, 89), our dataset reveals the extent and dynamics of these trade-offs on a genome-wide scale.

### Predicting function by association

Using clustering analysis, we identified several genes that lack a well-established function, and that clustered tightly with well-studied genes. We can use this co-clustering to suggest shared functions (90).

Temporal clustering analysis identified *Mtk-like, CG43920, IBIN* and *CG45045*, which shared similar expression dynamics with Imd-regulated AMPs (**Figure 6A**). Suggesting AMP-like functions to these less characterized genes can be supported by observations from literature: All four genes were previously found to respond to infection (47, 55), and *Mtk-like* and *CG43920* have been shown to encode small proteins predicted to be cationic (91), properties shared by known AMPs (92). *IBIN* and *CG45045* were also predicted to physically interact with antimicrobial peptide transcripts (91). On the other hand, Valanne *et al.* (46) found that *IBIN* overexpression increased levels of hemocytes and hemolymph glucose, suggesting that *IBIN* functions as a link between immunity and metabolism, instead of acting as an AMP itself.

The dense sampling nature of this time course allowed us to discern the clear cycling patterns of differentially expressed genes such as *period*, *timeless*, *takeout*, *vrille*, and *cryptochrome*, all of which have well-characterized circadian rhythm functions (38, 39, 76, 93, 94). Clustered with these genes, our analysis identified other genes that showcase cyclic behavior but are not canonically circadian-associated genes. This includes eight genes which do not have assigned circadian functions but do have some evidence of cyclic behavior in previous literature. It also includes ten genes that had not been reported to exhibit any cyclic expression before this study (**Table 1**). The identification of canonical circadian rhythm patterns both validates our methods of data normalization and differential expression analysis, and increases the certainty that we are accurately profiling novel temporal dynamics. It is important to note, however, that proper validation of the cycling behavior of our novel cyclic genes should be performed under normal *Drosophila* conditions, as we do not know whether Imd stimulation affected their expression.

Overall, we were able to implicate these uncharacterized genes as potential members of specific functional pathways due to the strong similarity of their expression dynamics. This is impactful both in the functional implication of these genes, but also in demonstrating the potential of this “guilt-by-association” method to assign putative function to other uncharacterized genes through RNA expression time-course experiments.

### Limitations and future steps

Limitations of our experimental design can be used as a guide to design future experiments. This time-course design lacks time-matched controls to account for expression changes associated with phenomena outside the Imd stimulation, such as aging. However, it is still highly valuable to develop and improve methods for analyzing time-course transcriptional data lacking time-matched controls, since these methods are needed to analyze processes such as development, where such controls are inherently not possible.

This study also lacks a control for the wounding injury caused by the injection itself. As discussed above, specific expression patterns such as those from the Toll-response genes could be caused by the injection wounding rather than the Gram-negative peptidoglycan in commercial LPS. Another important experimental design aspect for time series is choosing the time frame within which sampling needs to occur, which can be difficult to establish in advance. In our study Imd-regulated gene expression was sustained until five days after infection, and a more prolonged sampling would have been necessary to determine the full duration of these genes’ up-regulation after Imd stimulation.

Further, our experiment sampled only males, while several previously published immune response studies used females (e.g. (3, 21)). Overall, the observed gene expression changes and trade-offs in our study agreed with functional and omics studies done using females. This suggests that global gene expression dynamics after an Imd challenge are similar between males and females, but future experiments would be needed to make direct comparisons. In addition, multiple studies have reported a trade-off between immunity and reproduction, which occurs in singly-mated females (9) and multiply mated males (95). Mating activity was outside the scope of our design but can be included in future designs.

In our dataset, Granger causality analysis excelled at showcasing the relationships between divergent gene pairs, but was overly sensitive to the extreme temporal correlation between large groups of genes when analyzing positive edges. To avoid a prohibitively dense network for analysis, we relied on heuristic network trimming criteria, which was effective, but is likely not generalizable to other similar experiments. Developing co-integration methods that take into account the specific bias found in high-dimensional RNA-seq datasets would provide a more robust statistical analysis of the causal relationships observed in this type of data. The biological interpretation of relationships identified using Granger causality also warrants further investigation: Granger causality was successful at identifying what are likely the downstream results of divergent regulation, and it was successful at identifying positive lead-lag relationships between genes that likely respond to similar signals, but might differ in their exact (post-)transcriptional control. However, these statistical causal relationships provide only hypotheses that should be tested with direct experimental disruptions of a system to demonstrate biological causality.

## CONCLUSIONS

Our combination of analytical methods provides robust profiling of the innate immune response in *Drosophila melanogaster* after Imd stimulation at the highest temporal resolution to date, and serves as a proof of concept for high-density time-course RNA-seq analyses in other systems. Further, it motivates innovation in computational and statistical methods for longitudinal omics data, both to account for their inherent high-dimensionality and the complex underlying architecture that contains both causal and spurious coordination. Specifically, the development and application of multivariate Granger causality analysis highlights the potential of time-course data to evaluate coordinated gene expression changes through lags and trade-offs. While the immune response in *D. melanogaster* has been well studied, our research using dense time-course gene expression data reveals genome-wide dynamic expression patterns at higher temporal detail. Specifically, we reveal responses to Imd stimulation with divergent initiation and resolution dynamics, cyclic patterns of gene expression, and patterns of co-regulation and functional trade-offs, while also assigning putative gene functions to uncharacterized genes through a temporal guilt-by-association method.

## MATERIALS AND METHODS

### Fly lines, injections, and sample collection

Male adult *Drosophila* of about 4 days old were sampled from an F1 cross from two *Drosophila melanogaster* Genetic Reference Panel (DGRP) lines: line 379, which has shown to have low bacterial resistance, and line 360, which has high bacterial resistance (96). Offspring from these two DGRP lines were used to allow investigation of allele-specific expression after Imd stimulation. Analysis indicated significant allele-specific differential expression across multiple loci, including loci that contain immune genes, when aggregating data across all time points (data not shown). This may reflect differential rates of transcription initiation, processivity, RNA processing, or decay. However, allele-specific expression did not significantly change over time after infection (data not shown). Because of the absence of time-dependent effects, this analysis was not included in the manuscript.

Flies were kept on a 12:12 dark-light cycle, on standard yeast/glucose food. Flies were injected in the abdomen with 9.2 μl of commercial lipopolysaccharide (LPS) (*Escherichia coli* 055:B5 Sigma) derived from the outer membrane of Gram-negative bacteria. Flies were injected using a Nanoinjector (Nanoject II, catalog #3-000-204, Drummond), which allows high-throughput fly injections with a constant injection volume. Injections were performed in the abdomen, as it has been shown to be less detrimental to the fly compared to thorax injury (97).

Flies were sampled for a total of 21 time points throughout the course of five days, which included an uninfected un-injected baseline sample as control at time zero, and 20 time points after LPS injection (1, 2, 3, 4, 5, 6, 8, 10, 12, 14, 16, 20, 24, 30, 36, 42, 48, 72, 96, 120 hr). This sampling was performed twice, using flies from the same stock, on two consecutive days. Therefore, all time points have two replicates, giving a total of 42 samples. Since flies were sampled from the same stock for both replicates, we consider the second replicate to control for any effects of the injection technique. This sampling strategy was informed by experimental data and theoretical analysis that show that under reasonable assumptions, sampling time points at higher resolution is preferred over having more replicates (98), an important strategy to consider when having a limited experimental budget.

During collection, a group of ∼10 pooled flies corresponding to the sampled time point were flash frozen in dry ice and stored at −80 C for later RNA extraction.

### Experimental validation using qPCR

The Imd inducibility of commercial LPS was confirmed using qPCR. Adult male *Drosophila* were injected with 9.2 μl or 40 μl of 1 mg/mL LPS and flash frozen at 8 and 24 hr for RNA extraction. Uninfected un-injected flies were used as control. Each sampled time point consisted of a group of ∼10 pooled flies. Each sample had two replicates. Genes *AttA* and *DptB* were measured to confirm Imd stimulation. Gene *Rp49* was used as a baseline for expression normalization. Results showed a significant up-regulation of *AttA* and *DptB* at both volumes (9.2 μl) for both time points (8 and 24 hr). We decided to use 9.2 μl and 40 μl least amount of disruption to flies during infections, while still eliciting an immune response.

### RNA extraction, RNA sequencing, and quality control filtering

RNA extraction was performed using Trizol (Life Technologies) following the manufacturer’s instructions. cDNA libraries were prepared using the TruSeq RNA Sample Preparation Kit (Illumina). RNA purity was assessed using a Nanodrop instrument. RNA concentration was determined using a Qubit (Life Technologies) instrument. Sequencing was performed on an Illumina Hi-Seq 2500, single-end, and a read length of 75 bp, at Cornell Biotechnology Resource Center Genomics Facility.

Samples had an average of 24.8 M raw reads. Samples went through quality control using FastQC (v0.11.5). Truseq adapter sequences were removed from any sample that showed any level of adapter contamination using cutadapt (v1.14). Low quality bases in the beginning and end of the reads were trimmed using fastx_trimmer (v0.0.13, http://hannonlab.cshl.edu/fastx_toolkit/). Reads were mapped to the *Drosophila melanogaster* genome (r6.17) using STAR (v2.5.2b). BAM files were generated using SAMtools (v1.3.2). Only one sample (4B, at 3 hr) out of the original 42 failed to pass the quality thresholds, and all subsequent analysis used the remaining 41 samples. An average of 92.97% reads per library mapped uniquely to the *Drosophila melanogaster* genome. We ended up with an average of 23.4 million uniquely mapped reads per library.

Reads mapping to genes were counted using the R package *GenomicAlignments*. Genes with zero counts across all samples were removed (923 genes out of 17,736). Samples were normalized to library size. A “+1” count number was added to all genes before performing log_2_ transformation, to make sure values after transformation are finite, and stabilize the variance at low expression end. After normalization and log_2_ transformation, only genes with more than 5 counts in at least 2 samples were kept (removing 4,156 genes). We ended up with 12,657 genes for downstream analysis.

A heatmap of the row Z-scores of normalized counts for all 12,657 genes indicated that sample 6A (5 hr after commercial LPS injection) was an outlier for a subset of the 12,657 genes (**Figure S12A**), even though sample 6A did not appear as an outlier using Principal Component Analysis (see below and **Figure 1B**). We identified outlier genes in sample 6A by subtracting replicate A row Z-scores of normalized counts from replicate B. While for most time points the difference between samples A and B varied between −4 and 4, samples 6A and 6B demonstrated larger differences for 1,439 genes (**Figure S12B**). These genes were enriched for GO terms related to neuron signaling and development (based on PANTHER GO statistical overrepresentation test with FDR 0.05). Only 40 of the 1,439 genes were annotated as immune genes by Early *et al.* (31), and none of the 1,439 genes were among the DE genes detected as differentially expressed using pairwise comparisons or spline fitting. Library sizes for samples 6A and 6B were similar (26,256,507 for replicate A and 22,980,006 for replicate B). We think the difference between the replicates at 5 hr post-injection could be caused by unknown variation in the flies’ environment. It did not influence our data analysis and conclusions, but it is necessary to be aware of when using this dataset for other analyses.

### Principal component analysis

Principal components analysis (PCA) was performed using function *plotPCA* from the R package *DESeq2* (15) after regularized-logarithm transformation of raw counts, using the design ∼time+time:time to create the DEseqDataSet. Genes with zero counts across all samples were first removed. The default number of 500 top genes with highest row variance was used to calculate the principal components.

### Differential expression analysis

In order to identify genes that had differential expression over the time course, we adopted the linear model-based methodology proposed in Law *et al.* (18) and available in the R package *limma*. We first transformed the normalized RNA-seq read counts (before log_2_ transformation) using the *voom* transformation, which estimates the heteroscedastic mean variance relationships of log-counts and adds a precision weight to each observation to make them amenable to the usual linear modeling pipelines that rely on normality. We used gene-wise linear models to fit cubic splines (with 3 degrees of freedom) with time, TMM normalization method (99), and standard empirical Bayes *F*-tests to select genes whose expression levels were significantly altered across the time course in both replicates.

We also fit 3 degree polynomials across the first 48h using the R package maSigPro (19, 20). We used default parameters, including counts=F to model the data based on a normal distribution, since we ran maSigPro on counts that were normalized using Limma-Voom. We selected 169 genes as significantly time dependent if they had alfa (Benjamini-Hochberg corrected) < 0.05 and a goodness of fit R^2^ value of at least 0.6.

Next, we checked for differential expression of every gene between time point 0 (control) and time point *t*, for *t* = 1, 2, …, 48 hr. For each test, a multiple testing correction at 5% False Discovery Rate (FDR) using the Benjamini-Hochberg method (30) was adopted. Venn diagrams to compare results were adapted from those generated using web tool Venny (http://bioinfogp.cnb.csic.es/tools/venny/).

### Functional annotation

Gene Ontology (GO) enrichment analysis was performed using PANTHER Statistical Overrepresentation Test (http://pantherdb.org/, v14.1, released 2019/04/29) (100) using default settings (“GO biological process complete” as annotation dataset, Fisher’s Exact test, FDR < 0.05). Transcription factors were identified among differentially expressed genes based on a list of 753 putative site-specific transcription factors available via the FlyTF database (v1) (32). Gene set analysis was done using the R package *GSA*, which uses a Gene Set Analysis algorithm (101) that improves the GSEA algorithm (102) by allowing testing for associations between gene sets and time-dependent variables (101, 103). Gene set membership was assigned from GO data downloaded from FlyBase.org in January 2019 (version FB2019_01). Normalized counts for both replicates at each time point from 1 to 120 hr were compared against both control replicates (0 h), using a two-class paired vector (−1, 1, −2, 2) which corresponds to (control_replicateA, timepointX_replicateA, control_replicateB, timepointX_replicateB). We used 100,000 permutations to estimate false discovery rates. Only pathways with *P*-values below 0.05 and with 5 or more genes from our full dataset were kept. A subset of most relevant pathways was compiled by selecting pathways that had at least one gene from the subset of 551 most predominant time-dependent genes, and had a score of 2.5 or more in at least one time point from 1 to 48 hr. This gave us 41 unique pathways as shown in **Figure 9**.

### Cluster analyses

Hierarchical clustering of 91 core genes was performed using R package *hclust*, using Euclidean correlation as a distance metric. Hierarchical clustering of 551 predominant time-dependent genes was done using the default Pearson correlation with R package *pheatmap*. Cluster membership assignment and mean patterns of expression across time for genes within each cluster was performed as done in White *et al.* (26). Temporal clustering was performed using the R package *TSclust* (104). Normalized counts of both replicates were clustered using dissimilarity measures from Autocorrelation-based method (ACF), which computes the dissimilarity between two time series as the distance between their estimated simple autocorrelation coefficients (105). This method was used with a *P*-value cutoff of 0.05, and only top 1% correlation edges were further explored.

Cyclic gene patterns were identified using the JTK_Cycle algorithm (43) available in R package *JTK_Cycle*. Nine regularly distributed time points were subset from both replicates every 6 hr (0, 6, 12, 18, 24, 30, 36, 42, 48 hr). The time point corresponding to 18 hr was approximated by averaging normalized gene counts between time points 16 and 20 h. We looked for rhythms between 18-30 hr (4 to 6 time points) with a cutoff of BH *Q*-value < 0.05 and amplitude > 0.5.

### Network inference

Granger causality-based methods (22) were used to construct putative interaction networks among genes in the form of directed graphs with individual genes as nodes. A directed edge from gene *A* to gene *B* is added if the time course of gene *A* Granger-causes the time course of gene *B*. The notion of ‘Granger causality’ is popular in learning lead-lag relationships among two or more time series. Formally, if the time series of gene *A*, given by *x_t_*, has some power in predicting the expression of gene *B* at time *t* + 1, called *y_t+1,_* over and above *y_t_* and conditioned on an information set *l_t_*, then gene *A* is said to exert a *Granger causal* effect on gene *B*. Bivariate Granger causality uses a small information set *l_t_* = {*x_1:t_, y_1:t_*} and captures Granger causal relationship from gene *A* to gene *B* by testing whether the regression coefficient *β* in the following bivariate regression is different from zero:

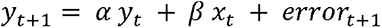

A master set of 258 genes was constructed from the 551 predominant time-dependent genes by picking those that had available functional annotation and that had differential expression of at least 2 log_2_ fold change. Using linear regression (function lm() in R), we conducted bivariate (pairwise) Granger causality tests for every pair of genes among this set of 258 genes using data on sliding windows of *t* = 6 consecutive time points and the two replicates (sample size = 12), and ranked them in order of increasing *P*-values (BH method used for calculating FDR), keeping the top resulting edges (BHFDR < 5%).

A well-known critique of bivariate Granger causality is its use of a small information set that does not contain any other factors except genes *A* and *B* (106). This failure to account for other potential confounding variables can give rise to many spurious edges in our network (106), where Granger causal effects from gene *A* to gene *B* is an artefact of gene *C*, which is causal for one or both genes. To address this, we adopted multivariate (or network) Granger causality (107), allowing us to avoid such spurious inferences through multiple linear regression. In this framework, we start with *p* genes, and Granger causal relationship of Gene *A* on Gene *B* is tested by regressing *y_t+1_* on *y_t_,x_t_* and the time courses of the other *p* - 2 genes *z*_1_*t*, *z*_2_*t, …, z_pt_*.

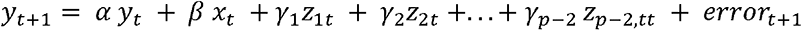

For small sample size and large *p*, the above regression is not possible to run using ordinary least squares (OLS), so we use LASSO (59) regression. To test if the regression coefficient *β* in the above regression is different from zero, we used two different variants of de-biased LASSO (60, 61), each of which corrects the bias of lasso and allows quantifying uncertainty of regression coefficients one at a time. A non-zero coefficient in the above multivariate regression suggests that gene *A* is Granger causal for gene *B*, even after accounting for the effects of the other *p*-2 genes. Using this method on the master set of 258 genes, we reconstructed putative directed networks of multivariate Granger causality and ranked the edges in increasing order of *P*-values, following the same parameters used in the bivariate (pairwise) Granger causality method (sliding window of 6 consecutive time points in both replicates, keeping the top resulting edges (BHFDR < 5%)).

## DECLARATIONS

### Ethics approval and consent to participate

Not applicable

### Consent for publication

Not applicable

### Availability of data and materials

The RNA-seq data will be available at NCBI SRA, BioProject ID PRJNA641552. The pipeline scripts and intermediary data files will be made available through GitHub. A query interface will allow any user to query a gene of interest and download a table of raw counts, normalized counts, log2 fold change, and plots of gene trajectory over time.

### Authors’ contributions

FS, AE, and AGC conceived the study. FS collected the samples and generated the time course data. FS, SB, MTW, and AGC conceived the computational and statistical analyses. FS, SYND, and SB performed the computational and statistical analyses. FS, SYND, SB, and AGC wrote the manuscript. All authors read and approved the final manuscript.

## Acknowledgements

The authors want to thank Juan Felipe Beltrán, Amanda Manfredo, Keegan Kelsey, Yasir Ahmed-Braimah, and Elissa Cosgrove for all the help and advice provided. We also thank the three anonymous reviewers for their comments and suggestions.

## SUPPORTING INFORMATION

### SUPPLEMENTAL FIGURES

**Figure S1.**
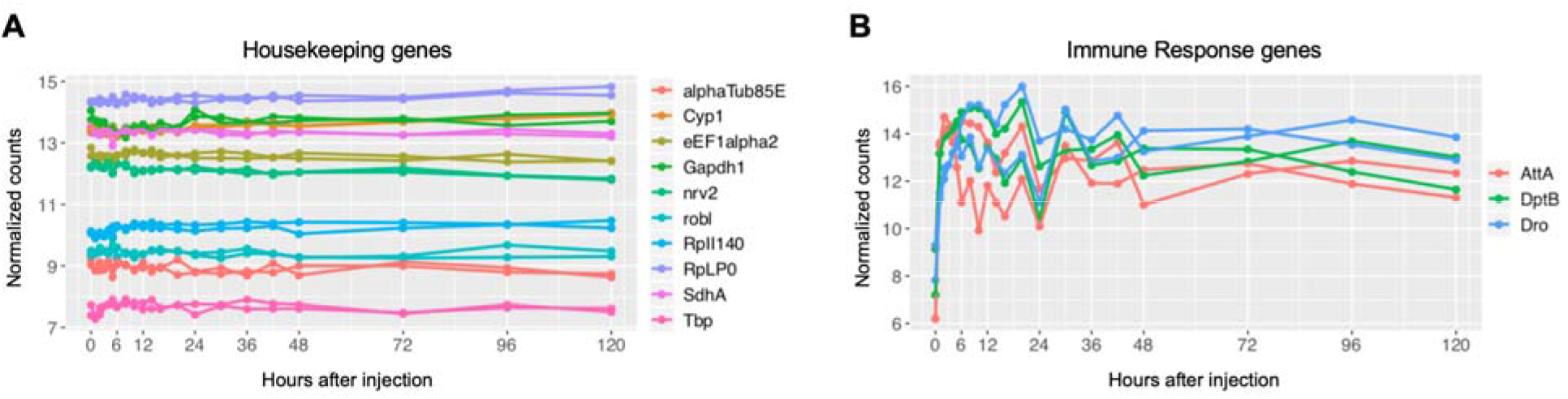
Plots of normalized counts of housekeeping genes (**A**) show little change across time as expected under proper data normalization, while immune response genes (**B**) show up-regulation within the first time points, as expected after a successful Imd stimulation.

**Figure S2.**
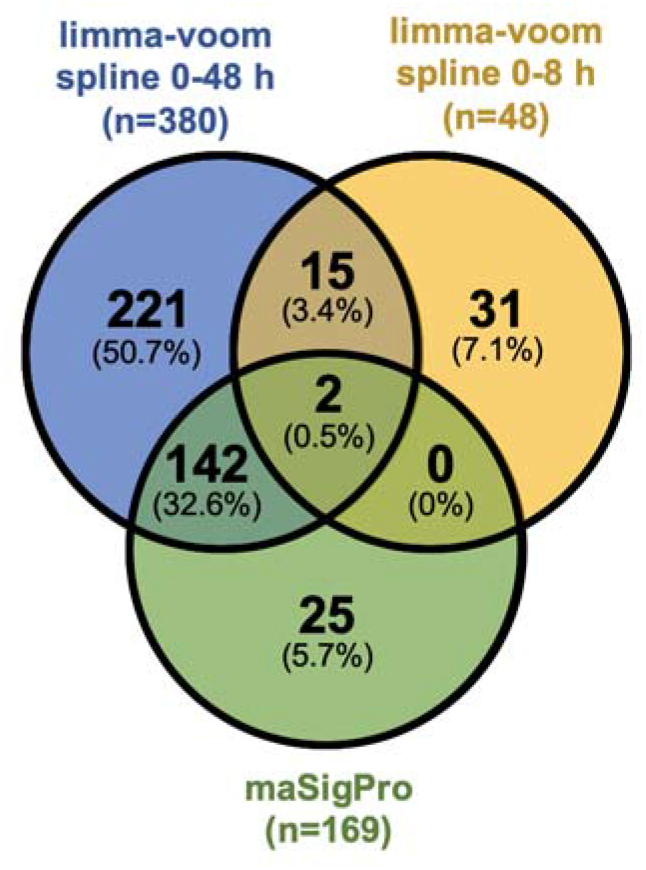
Venn Diagram showing overlap and differences of DE genes identified using limma-voom spline fitting vs. maSigPro fitting of polynomials.

**Figure S3.**
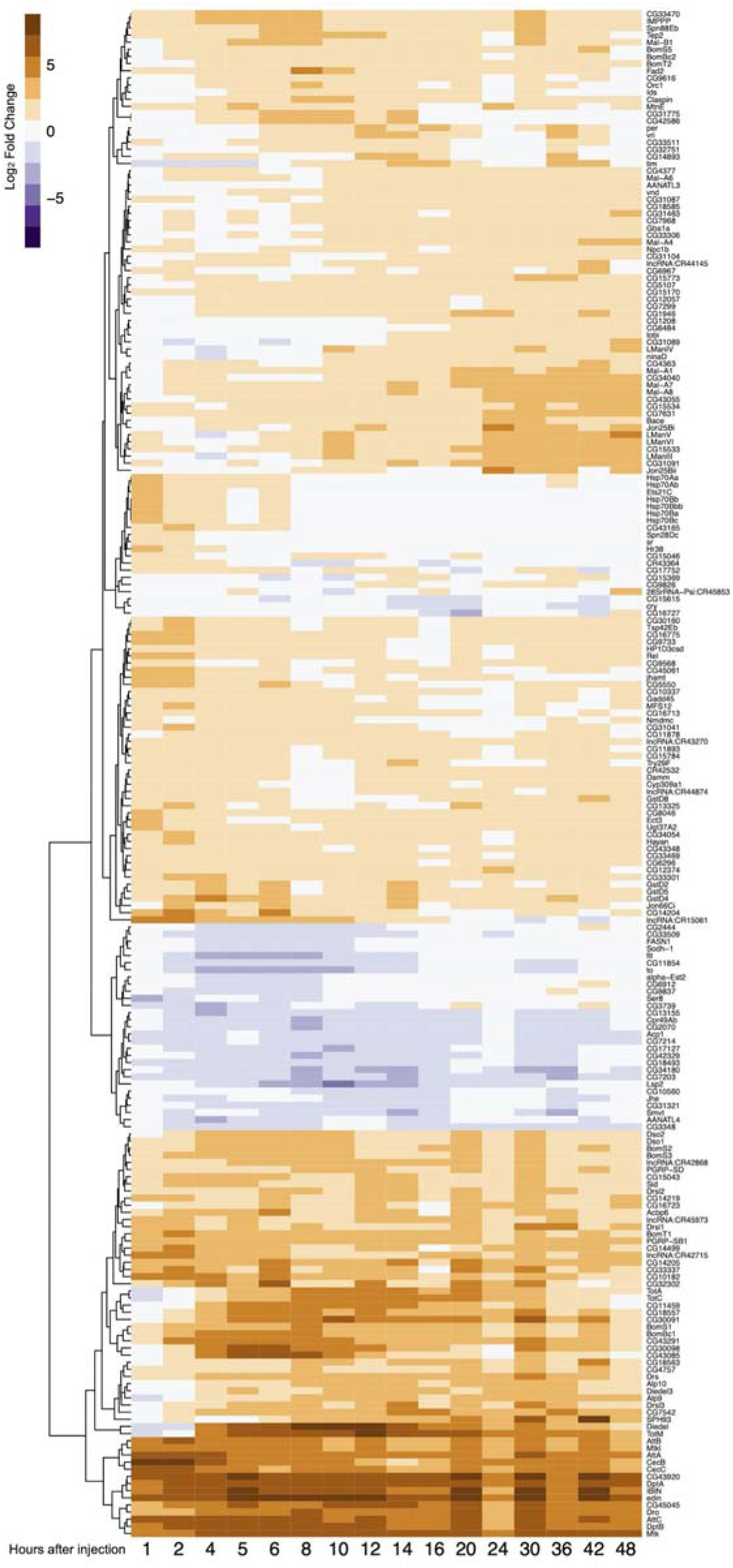
Heatmap of 214 genes differentially expressed following injection with commercial LPS. Scale is log_2_ fold change, with negative expression in purple and positive expression in dark orange, compared to baseline.

**Figure S4.**
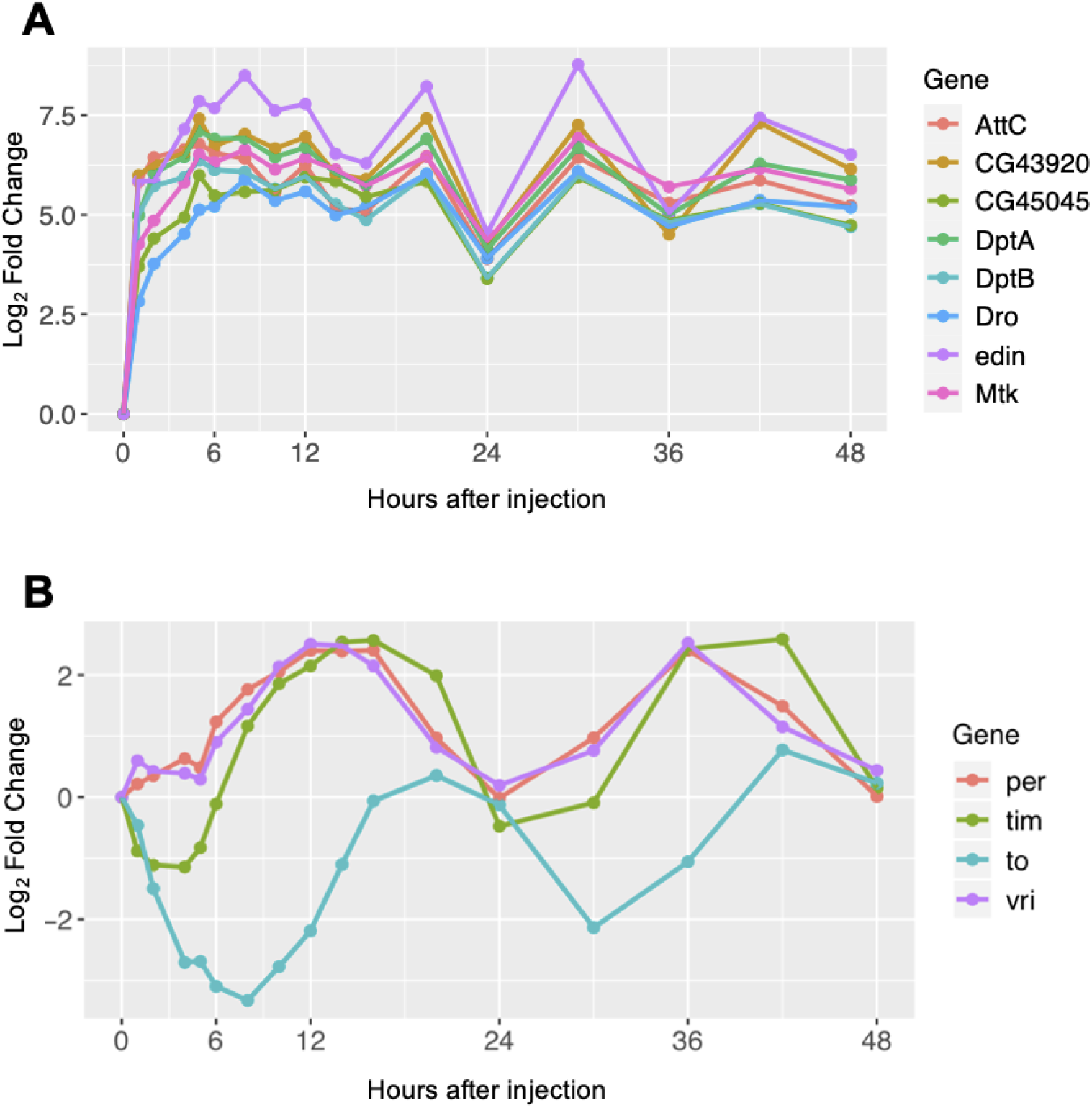
**(A)** Temporal dynamics of the most strongly up-regulated genes (*DptB*, *AttC*, *Mtk*, *Dro*, *CG45045*, *DptA*, *CG43920*, and *edin*) in the first 48 hr after injection. These genes are part of the 91 “core” DE genes with log_2_FC > 2 in at least two time intervals after injection, with an FDR < 0.01 (Figure 3). (**B**) Temporal dynamics of gene expression of circadian rhythm genes (*per*, *tim*, *to*, *vri*) show a classic and well characterized 24 hr periodic expression.

**Figure S5.**
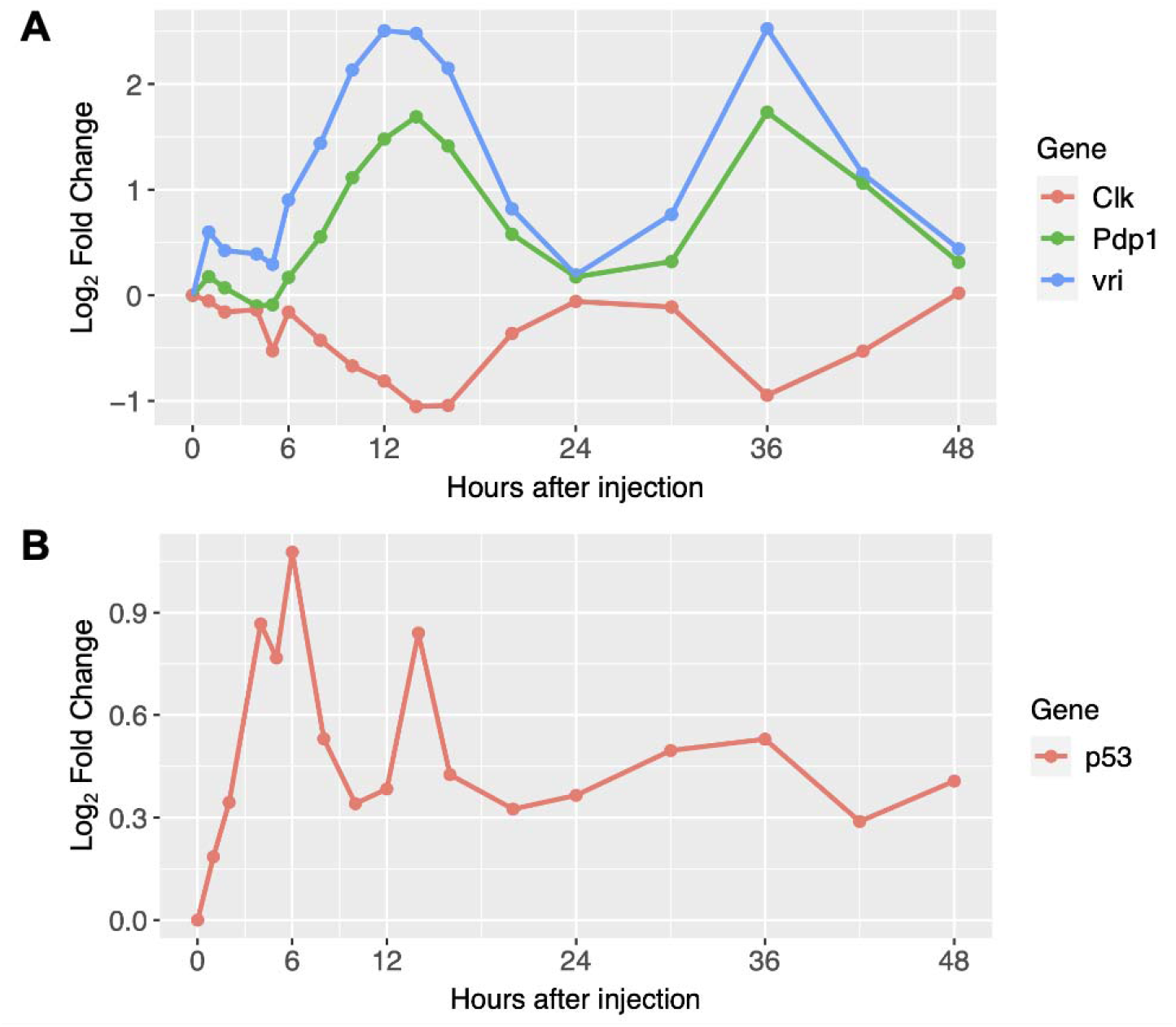
Expression profiles of DE genes encoding transcription factors. **(A)** Genes involved in regulation of the circadian clock. **(B)** *p53*, which encodes a transcription factor involved in DNA repair and genotoxic stress.

**Figure S6.**
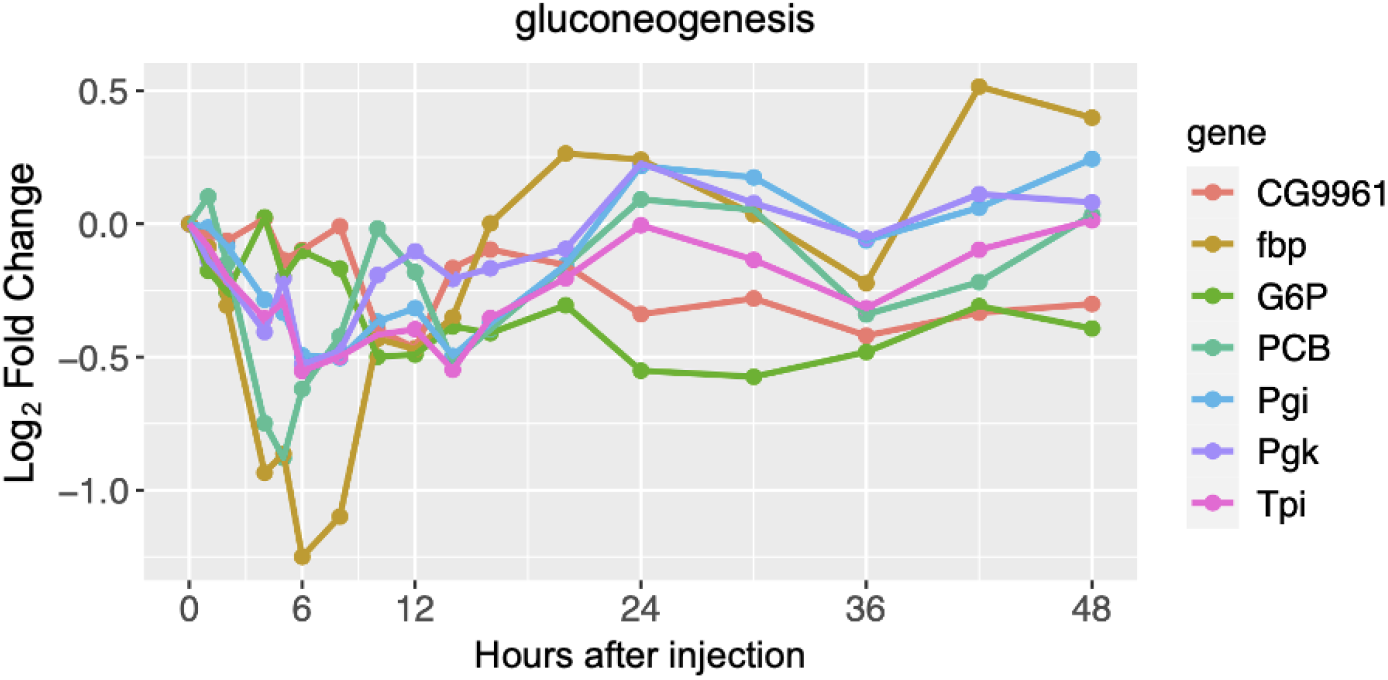
Gluconeogenesis pathway

**Figure S7.**
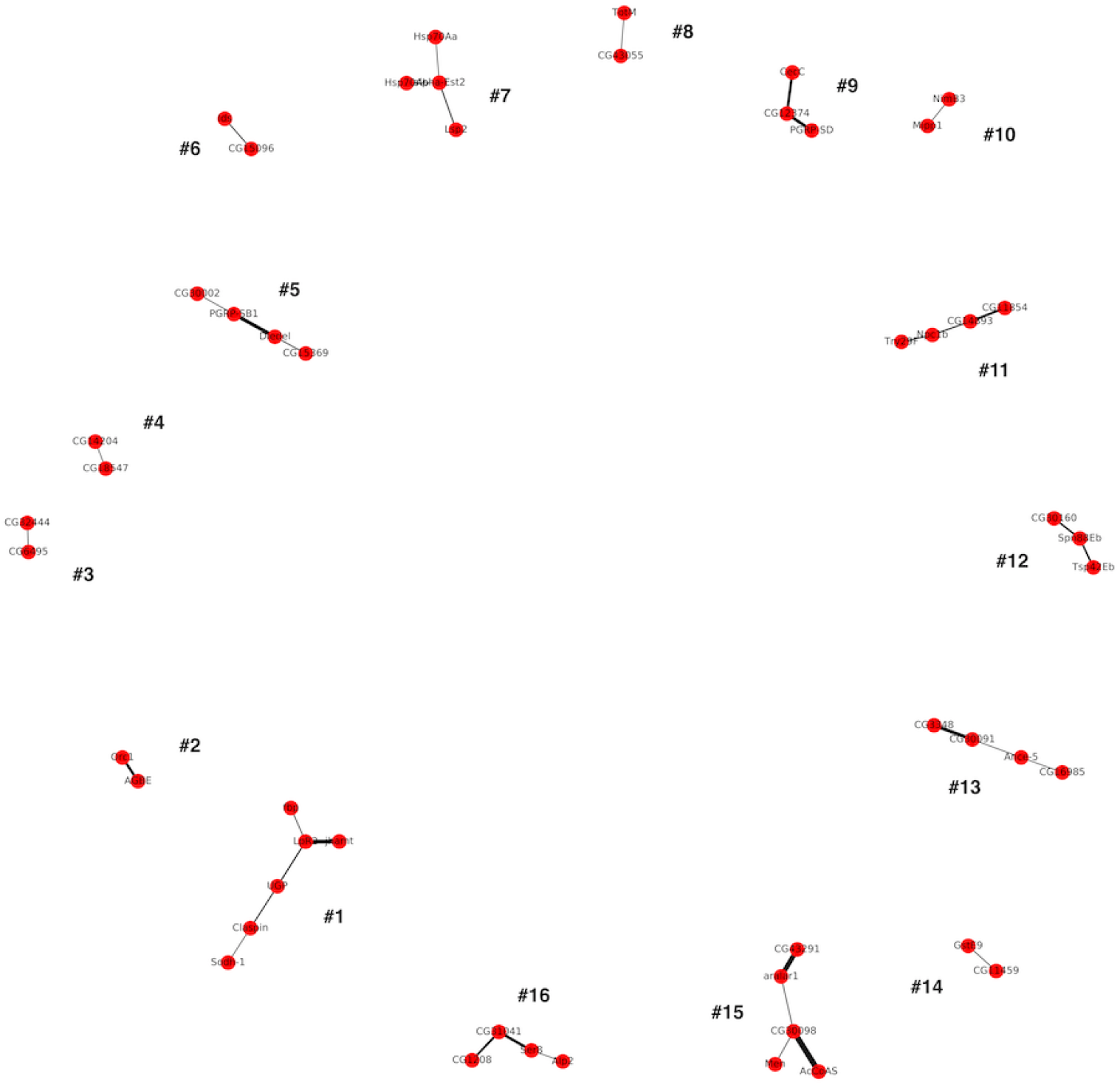
GC filtered network. The network was pruned to only the edges with at least three consecutive windows of GC significance, those that have a negative GC relationship, and it excludes genes identified with JTK_Cycle.

**Figure S8.**
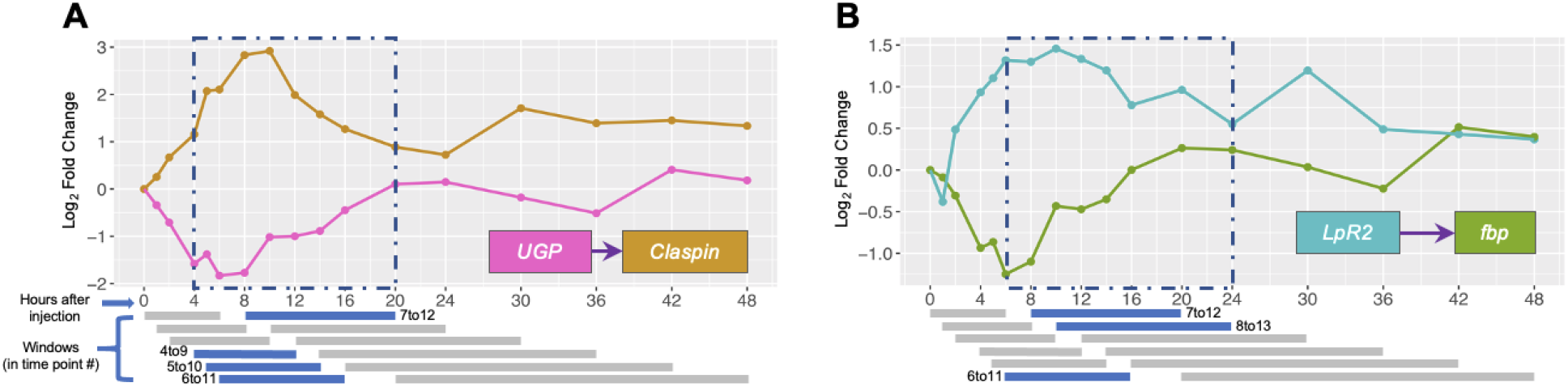
Negative GC edges. **(A) between *Claspin* and *UGP.* (B) between *LpR2* and *fbp*** Windows in which a significant Granger Causal relationship is established are colored in blue and are contained in a dashed blue box marking the plot area, non-significant windows are grey below the plot.

**Figure S9.**
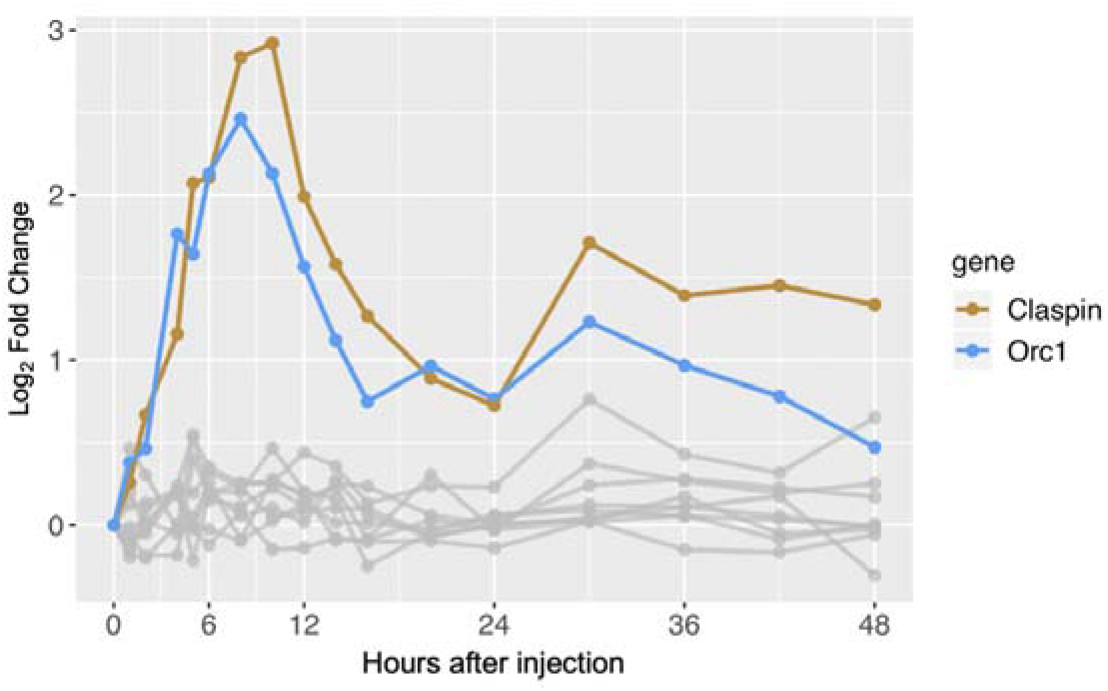
Pathway corresponding to ‘mitotic DNA replication checkpoint’.

**Figure S10.**
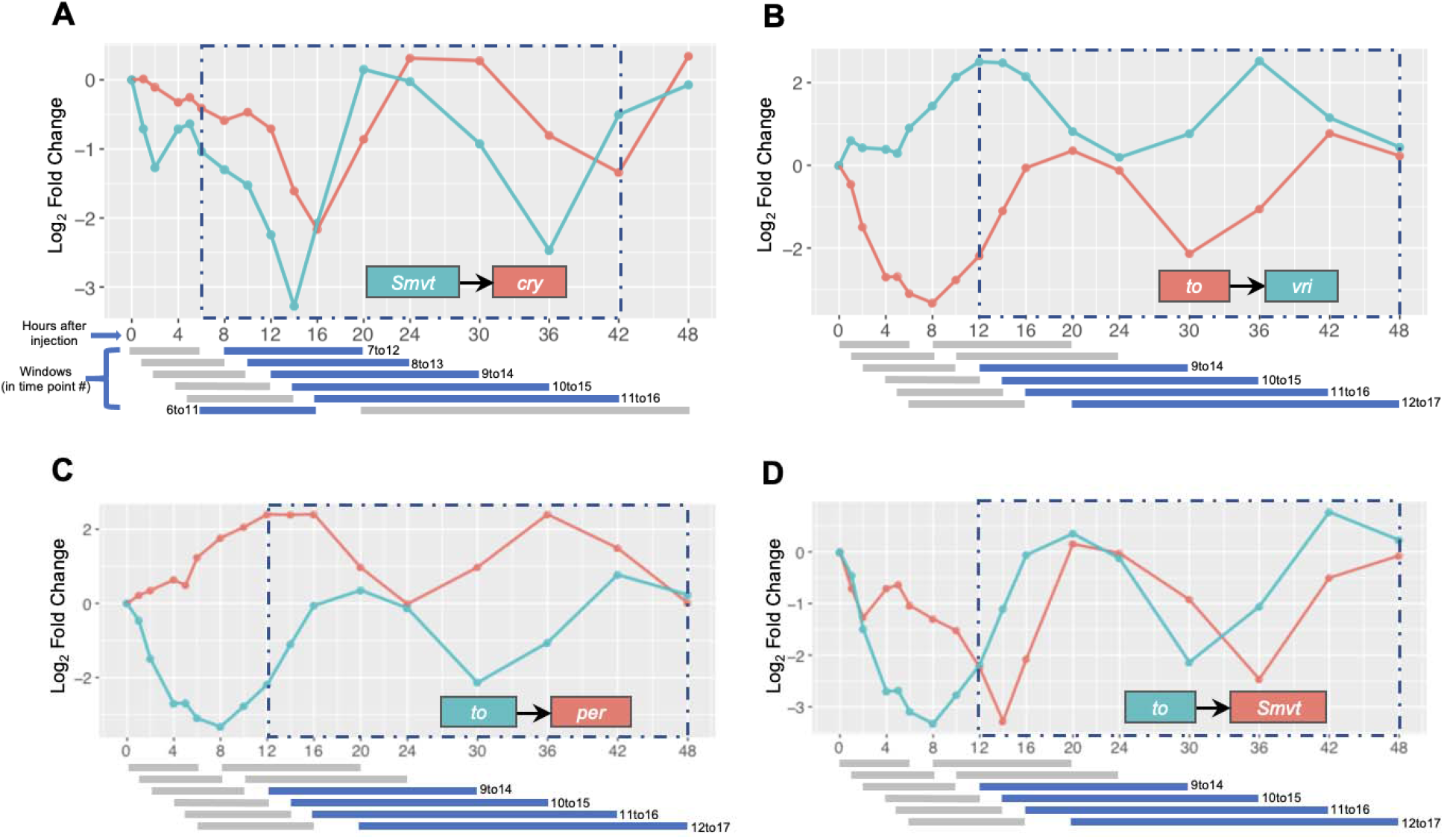
GC edges of circadian rhythm genes plotted against time. (**A**) Positive lead-lag correlation between *Smvt* and *cry*. (**B**) Negative lead-lag correlation between *to* and *vri*. (**C**) Negative lead-lag correlation between *period* and *takeout.* (**D**) Positive lead-lag correlation between *Smvt* and *takeout*. Significant windows are colored in blue, non-significant windows are colored in grey. Resulting overall consecutive windows are labeled in blue dashed rectangles. Individual windows represent 6 consecutive time points. Note that time points are not regularly distributed with time, therefore windows have different time ranges, but identical number of samples.

**Figure S11.**
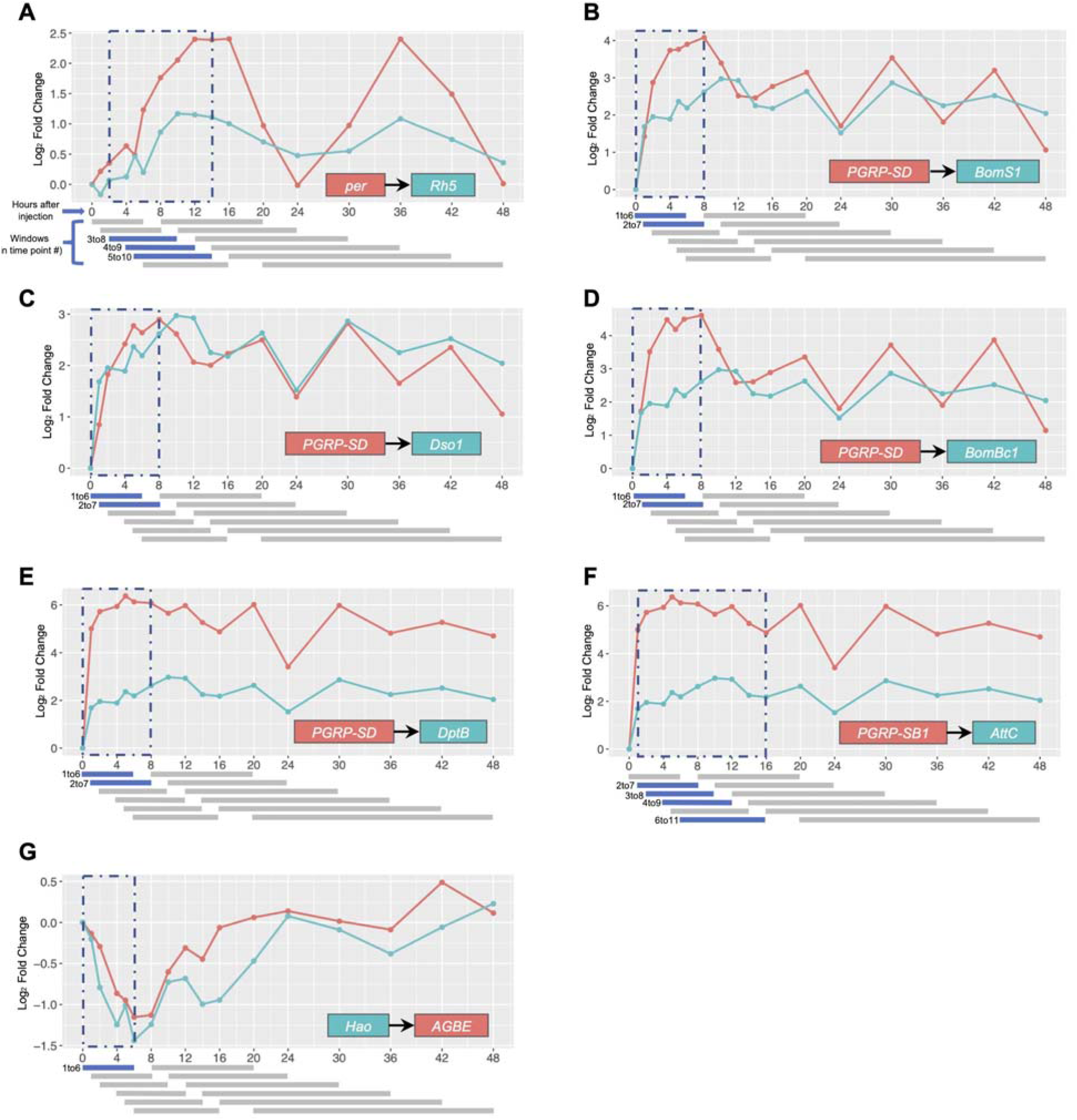
Positive GC edges. Positive lead-lag correlation between (**A**) *period* and *Rh5*, (**B**) *BomS1* and *PGRP-SD*, (**C**) *Dso1* and *PGRP-SD*, (**D**) *BomBc1* and *PGRP-SD*, (**E**) *DptB* and *PGRP-SD*, (**F**) *AttC* and *PGRP-SB1*, and (**G**) *AGBE* and *Hao.* Significant windows are colored in blue, non-significant windows are colored in grey. Resulting overall consecutive windows are labeled in blue dashed rectangles. Individual windows represent 6 consecutive time points. Note that time points are not regularly distributed with time, therefore windows have different time ranges, but identical number of samples.

**Figure S12.**
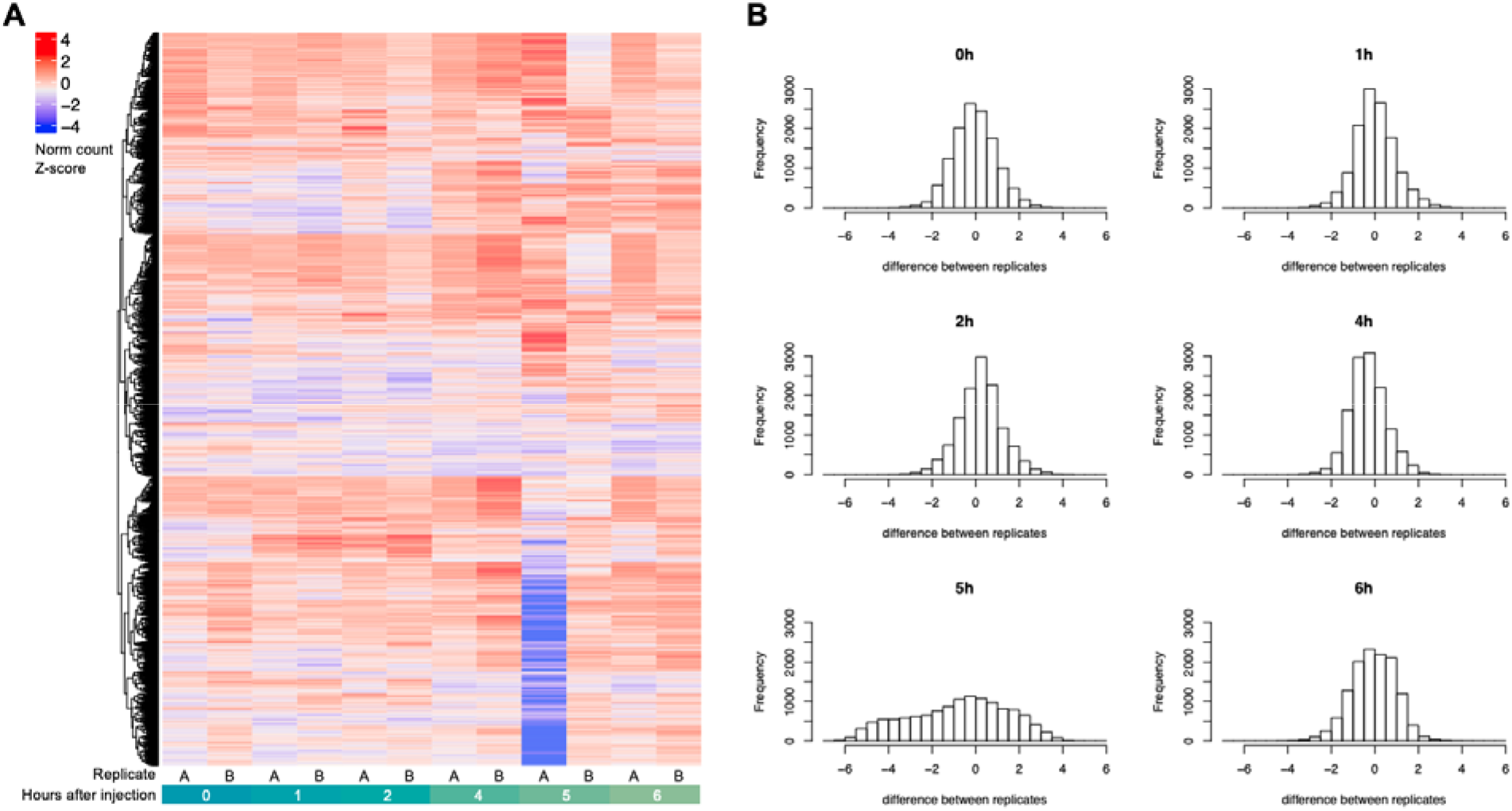
Sample 6A is an outlier for a subset of genes. **A**) Heatmap showing row Z scores of normalized counts for all 12,657 genes in the filtered datasets. Each row represents a gene; each column represents a replicate, from pre-injection (0 hours) up to 6 hr post-injection. (**B**) Distribution of differences between normalized count Z scores for replicates A and B for each time point.

**Table S1.** List of differentially expressed genes identified using spline fitting and pairwise.

**Table S2.**
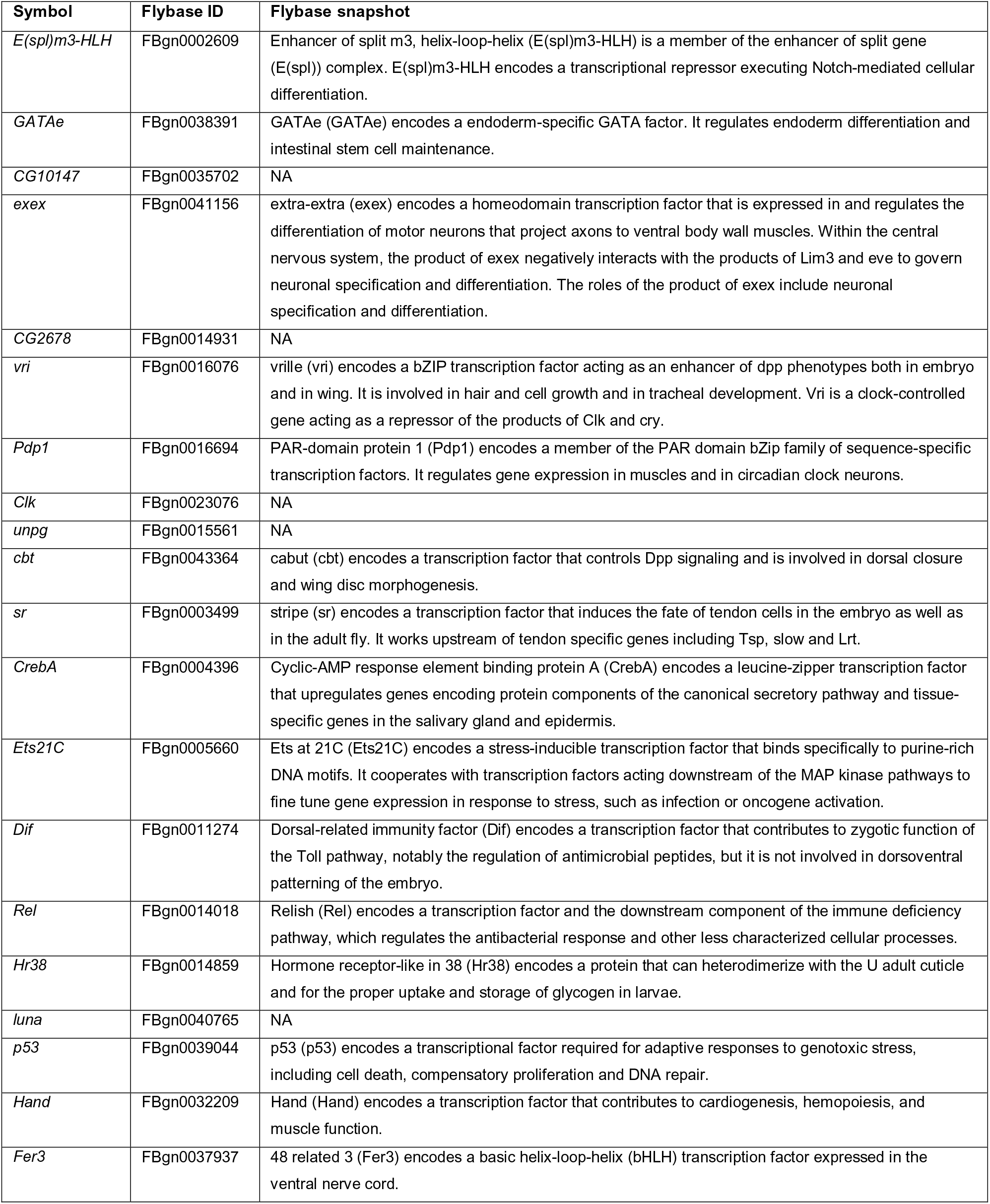
20 genes that respond to commercial LPS injection encode transcription factors.

## REFERENCES

1. Lazzaro BP, Galac Madeline R. Disease Pathology: Wasting Energy Fighting Infection. Current Biology. 2006;16(22):R964–R5.

2. Zerofsky M, Harel E, Silverman N, Tatar M. Aging of the innate immune response in *Drosophila melanogaster*. Aging Cell. 2005;4(2):103–8.

3. DiAngelo JR, Bland ML, Bambina S, Cherry S, Birnbaum MJ. The immune response attenuates growth and nutrient storage in Drosophila by reducing insulin signaling. Proceedings of the National Academy of Sciences of the United States of America. 2009;106(49):20853–8.

4. Fitzpatrick M, Young SP. Metabolomics – A novel window into inflammatory disease. Swiss medical weekly. 2013;143:w13743-w.

5. McKean KA, Yourth CP, Lazzaro BP, Clark AG. The evolutionary costs of immunological maintenance and deployment. BMC Evolutionary Biology. 2008;8(1):76.

6. Howick VM, Lazzaro BP. Genotype and diet shape resistance and tolerance across distinct phases of bacterial infection. BMC Evolutionary Biology. 2014;14(1):56.

7. Fedorka KM, Linder JE, Winterhalter W, Promislow D. Post-mating disparity between potential and realized immune response in Drosophila melanogaster. Proceedings of the Royal Society B: Biological Sciences. 2007;274(1614):1211–7.

8. Short SM, Lazzaro BP. Female and male genetic contributions to post-mating immune defence in female Drosophila melanogaster. Proceedings of the Royal Society B: Biological Sciences. 2010;277(1700):3649–57.

9. Short SM, Wolfner MF, Lazzaro BP. Female *Drosophila melanogaster* suffer reduced defense against infection due to seminal fluid components. Journal of insect physiology. 2012;58(9):1192–201.

10. Schwenke RA, Lazzaro BP, Wolfner MF. Reproduction–Immunity Trade-Offs in Insects. Annual review of entomology. 2016;61:239–56.

11. Boutros M, Agaisse H, Perrimon N. Sequential Activation of Signaling Pathways during Innate Immune Responses in Drosophila. Developmental Cell. 2002;3(5):711–22.

12. De Gregorio E, Spellman PT, Rubin GM, Lemaitre B. Genome-wide analysis of the Drosophila immune response by using oligonucleotide microarrays. Proceedings of the National Academy of Sciences of the United States of America. 2001;98(22):12590–5.

13. Sackton TB, Lazzaro BP, Clark AG. Genotype and Gene Expression Associations with Immune Function in Drosophila. PLOS Genetics. 2010;6(1):e1000797.

14. Robinson MD, McCarthy DJ, Smyth GK. edgeR: a Bioconductor package for differential expression analysis of digital gene expression data. Bioinformatics. 2010;26(1):139–40.

15. Love MI, Huber W, Anders S. Moderated estimation of fold change and dispersion for RNA-seq data with DESeq2. Genome Biology. 2014;15(12):550.

16. Bar-Joseph Z, Gitter A, Simon I. Studying and modelling dynamic biological processes using time-series gene expression data. Nature Reviews Genetics. 2012;13:552.

17. Spies D, Ciaudo C. Dynamics in Transcriptomics: Advancements in RNA-seq Time Course and Downstream Analysis. Computational and Structural Biotechnology Journal. 2015;13:469–77.

18. Law CW, Chen Y, Shi W, Smyth GKJGB. voom: precision weights unlock linear model analysis tools for RNA-seq read counts. Genome Biology. 2014;15(2):R29.

19. Conesa A, Nueda MJ, Ferrer A, Talón M. maSigPro: a method to identify significantly differential expression profiles in time-course microarray experiments. Bioinformatics. 2006;22(9):1096–102.

20. Nueda MJ, Tarazona S, Conesa A. Next maSigPro: updating maSigPro bioconductor package for RNA-seq time series. Bioinformatics. 2014;30(18):2598–602.

21. Kaneko T, Goldman WE, Mellroth P, Steiner H, Fukase K, Kusumoto S, et al. Monomeric and Polymeric Gram-Negative Peptidoglycan but Not Purified LPS Stimulate the Drosophila IMD Pathway. Immunity. 2004;20(5):637–49.

22. Granger CWJ. Investigating causal relations by econometric models and cross-spectral methods. Econometrica. 1969;37(3):424–38.

23. Graham AL, Shuker DM, Pollitt LC, Auld SKJR, Wilson AJ, Little TJ. Fitness consequences of immune responses: strengthening the empirical framework for ecoimmunology. Functional Ecology. 2011;25(1):5–17.

24. Deng Q, Ramsköld D, Reinius B, Sandberg R. Single-Cell RNA-Seq Reveals Dynamic, Random Monoallelic Gene Expression in Mammalian Cells. Science. 2014;343(6167):193–6.

25. Bendjilali N, MacLeon S, Kalra G, Willis SD, Hossian AKMN, Avery E, et al. Time-Course Analysis of Gene Expression During the Saccharomyces cerevisiae Hypoxic Response. G3: Genes|Genomes|Genetics. 2017;7(1):221–31.

26. White RJ, Collins JE, Sealy IM, Wali N, Dooley CM, Digby Z, et al. A high-resolution mRNA expression time course of embryonic development in zebrafish. eLife. 2017;6:e30860.

27. Novembre J, Stephens M. Interpreting principal component analyses of spatial population genetic variation. Nature genetics. 2008;40(5):646–9.

28. Podani J, Miklós I. Resemblance Coefficients and The Horseshoe Effect in Principal Coordinates Analysis. Ecology. 2002;83(12):3331–43.

29. Lü J, Yang C, Zhang Y, Pan H. Selection of Reference Genes for the Normalization of RT-qPCR Data in Gene Expression Studies in Insects: A Systematic Review. Frontiers in Physiology. 2018;9:1560-.

30. Benjamini Y, Hochberg Y. Controlling the False Discovery Rate: A Practical and Powerful Approach to Multiple Testing. Journal of the Royal Statistical Society Series B (Methodological). 1995;57(1):289–300.

31. Early AM, Arguello JR, Cardoso-Moreira M, Gottipati S, Grenier JK, Clark AG. Survey of Global Genetic Diversity Within the Drosophila Immune System. Genetics. 2017;205(1):353.

32. Pfreundt U, James DP, Tweedie S, Wilson D, Teichmann SA, Adryan B. FlyTF: improved annotation and enhanced functionality of the *Drosophila* transcription factor database. Nucleic acids research. 2010;38(Database issue):D443–D7.

33. Meng X, Khanuja BS, Ip YT. Toll receptor-mediated Drosophila immune response requires Dif, an NF-κB factor. Genes & Development. 1999;13(7):792–7.

34. Manfruelli P, Reichhart J-M, Steward R, Hoffmann JA, Lemaitre B. A mosaic analysis in *Drosophila* fat body cells of the control of antimicrobial peptide genes by the Rel proteins Dorsal and DIF. The EMBO Journal. 1999;18(12):3380–91.

35. Myllymäki H, Valanne S, Rämet M. The *Drosophila* Imd Signaling Pathway. The Journal of Immunology. 2014;192(8):3455.

36. Mundorf J, Donohoe CD, McClure CD, Southall TD, Uhlirova M. Ets21c Governs Tissue Renewal, Stress Tolerance, and Aging in the *Drosophila* Intestine. Cell Reports. 2019;27(10):3019–33.e5.

37. Chen X, Rahman R, Guo F, Rosbash M. Genome-wide identification of neuronal activity-regulated genes in Drosophila. eLife. 2016;5:e19942.

38. Cyran SA, Buchsbaum AM, Reddy KL, Lin MC, Glossop NR, Hardin PE, et al. *vrille*, *Pdp1*, and *dClock* form a second feedback loop in the *Drosophila* circadian clock. Cell. 2003;112(3):329–41.

39. Collins B, Mazzoni EO, Stanewsky R, Blau J. *Drosophila* CRYPTOCHROME Is a Circadian Transcriptional Repressor. Current Biology. 2006;16(5):441–9.

40. Brodsky MH, Weinert BT, Tsang G, Rong YS, McGinnis NM, Golic KG, et al. *Drosophila melanogaster* MNK/Chk2 and p53 Regulate Multiple DNA Repair and Apoptotic Pathways following DNA Damage. Molecular and Cellular Biology. 2004;24(3):1219–31.

41. Hoffmann JA, Reichhart J-M. Drosophila innate immunity: an evolutionary perspective. Nature Immunology. 2002;3:121.

42. Miyamoto T, Amrein H. Gluconeogenesis: An ancient biochemical pathway with a new twist. Fly (Austin). 2017;11(3):218–23.

43. Hughes ME, Hogenesch JB, Kornacker K. JTK_CYCLE: an efficient nonparametric algorithm for detecting rhythmic components in genome-scale data sets. J Biol Rhythms. 2010;25(5):372–80.

44. Cirelli C, LaVaute TM, Tononi G. Sleep and wakefulness modulate gene expression in *Drosophila*. Journal of Neurochemistry. 2005;94(5):1411–9.

45. Shirasu-Hiza MM, Dionne MS, Pham LN, Ayres JS, Schneider DS. Interactions between circadian rhythm and immunity in *Drosophila melanogaster*. Current Biology. 2007;17(10):R353–R5.

46. Valanne S, Salminen TS, Järvelä-Stölting M, Vesala L, Rämet M. Correction: Immune-inducible non-coding RNA molecule *lincRNA-IBIN* connects immunity and metabolism in *Drosophila melanogaster*. PLOS Pathogens. 2019;15(10):e1008088.

47. Troha K, Im JH, Revah J, Lazzaro BP, Buchon N. Comparative transcriptomics reveals CrebA as a novel regulator of infection tolerance in D. melanogaster. PLoS Pathogens. 2018;14(2):e1006847.

48. Clemmons AW, Lindsay SA, Wasserman SA. An Effector Peptide Family Required for *Drosophila* Toll-Mediated Immunity. PLOS Pathogens. 2015;11(4):e1004876.

49. Cohen LB, Lindsay SA, Xu Y, Lin SJH, Wasserman SA. The Daisho Peptides Mediate *Drosophila* Defense Against a Subset of Filamentous Fungi. Frontiers in Immunology. 2020;11(9).

50. Kenmoku H, Hori A, Kuraishi T, Kurata S. A novel mode of induction of the humoral innate immune response in *Drosophila* larvae. Disease Models & Mechanisms. 2017;10(3):271.

51. Ekengren S, Tryselius Y, Dushay MS, Liu G, Steiner H, Hultmark D. A humoral stress response in Drosophila. Current Biology. 2001;11(9):714–8.

52. Ao J, Ling E, Yu X-Q. Drosophila C-type lectins enhance cellular encapsulation. Mol Immunol. 2007;44(10):2541–8.

53. Keebaugh ES, Schlenke TA. Adaptive evolution of a novel *Drosophila* lectin induced by parasitic wasp attack. Mol Biol Evol. 2012;29(2):565–77.

54. Zsámboki J, Csordás G, Honti V, Pintér L, Bajusz I, Galgóczy L, et al. *Drosophila* Nimrod proteins bind bacteria. Central European Journal of Biology. 2013;8(7):633–45.

55. Katzenberger RJ, Ganetzky B, Wassarman DA. Age and Diet Affect Genetically Separable Secondary Injuries that Cause Acute Mortality Following Traumatic Brain Injury in Drosophila. G3: Genes|Genomes|Genetics. 2016;6(12):4151–66.

56. Buchon N, Poidevin M, Kwon HM, Guillou A, Sottas V, Lee BL, et al. A single modular serine protease integrates signals from pattern-recognition receptors upstream of the *Drosophila* Toll pathway. Proc Natl Acad Sci U S A. 2009;106(30):12442–7.

57. Fujita A, Severino P, Kojima K, Sato JR, Patriota AG, Miyano S. Functional clustering of time series gene expression data by Granger causality. BMC Systems Biology. 2012;6(1):137.

58. Finkle JD, Wu JJ, Bagheri N. Windowed Granger causal inference strategy improves discovery of gene regulatory networks. Proceedings of the National Academy of Sciences. 2018;115(9):2252.

59. Tibshirani R. Regression Shrinkage and Selection via the Lasso. Journal of the Royal Statistical Society Series B (Methodological). 1996;58(1):267–88.

60. Javanmard A, Montanari A. Confidence intervals and hypothesis testing for high-dimensional regression. J Mach Learn Res. 2014;15(1):2869–909.

61. Dezeure R, Buhlmann P, Meier L, Meinshausen N. High-Dimensional Inference: Confidence Intervals, p-Values and R-Software hdi. Statist Sci. 2015;30(4):533–58.

62. Lee E-M, Trinh TTB, Shim HJ, Park S-Y, Nguyen TTT, Kim M-J, et al. Drosophila Claspin is required for the G2 arrest that is induced by DNA replication stress but not by DNA double-strand breaks. DNA Repair. 2012;11(9):741–52.

63. Ubhi T, Brown GW. Exploiting DNA Replication Stress for Cancer Treatment. Cancer Research. 2019.

64. Liu D, Shaukat Z, Saint RB, Gregory SL. Chromosomal instability triggers cell death via local signalling through the innate immune receptor Toll. Oncotarget. 2015;6(36):38552–65.

65. Nakad R, Schumacher B. DNA Damage Response and Immune Defense: Links and Mechanisms. Front Genet. 2016;7:147-.

66. Soukup SF, Culi J, Gubb D. Uptake of the necrotic serpin in Drosophila melanogaster via the lipophorin receptor-1. PLoS genetics. 2009;5(6):e1000532-e.

67. Karlsson C, Korayem AM, Scherfer C, Loseva O, Dushay MS, Theopold U. Proteomic Analysis of the Drosophila Larval Hemolymph Clot. Journal of Biological Chemistry. 2004;279(50):52033–41.

68. Krautz R, Arefin B, Theopold U. Damage signals in the insect immune response. Front Plant Sci. 2014;5:342-.

69. Shinoda T, Itoyama K. Juvenile hormone acid methyltransferase: A key regulatory enzyme for insect metamorphosis. Proceedings of the National Academy of Sciences of the United States of America. 2003;100(21):11986–91.

70. Rolff J, Siva-Jothy MT. Copulation corrupts immunity: A mechanism for a cost of mating in insects. Proceedings of the National Academy of Sciences of the United States of America. 2002;99(15):9916–8.

71. Flatt T, Heyland A, Rus F, Porpiglia E, Sherlock C, Yamamoto R, et al. Hormonal Regulation of the Humoral Innate Immune Response in Drosophila melanogaster. The Journal of experimental biology. 2008;211(Pt 16):2712–24.

72. Schwenke RA, Lazzaro BP. Juvenile Hormone Suppresses Resistance to Infection in Mated Female Drosophila melanogaster. Current Biology. 2017;27(4):596–601.

73. Sarov-Blat L, So WV, Liu L, Rosbash M. The *Drosophila takeout* Gene Is a Novel Molecular Link between Circadian Rhythms and Feeding Behavior. Cell. 2000;101(6):647–56.

74. Wong R, Piper MDW, Wertheim B, Partridge L. Quantification of Food Intake in *Drosophila*. PLOS ONE. 2009;4(6):e6063.

75. Giebultowicz JM. Circadian regulation of metabolism and healthspan in *Drosophila*. Free Radical Biology and Medicine. 2018;119:62–8.

76. So WV, Sarov-Blat L, Kotarski CK, McDonald MJ, Allada R, Rosbash M. *takeout*, a Novel *Drosophila* Gene under Circadian Clock Transcriptional Regulation. Molecular and Cellular Biology. 2000;20(18):6935.

77. Smith RF, Konopka RJ. Effects of dosage alterations at the *per* locus on the period of the circadian clock of *Drosophila*. Molecular and General Genetics MGG. 1982;185(1):30–6.

78. Reddy P, Zehring WA, Wheeler DA, Pirrotta V, Hadfield C, Hall JC, et al. Molecular analysis of the *period* locus in *Drosophila melanogaster* and identification of a transcript involved in biological rhythms. Cell. 1984;38(3):701–10.

79. Claridge-Chang A, Wijnen H, Naef F, Boothroyd C, Rajewsky N, Young MW. Circadian Regulation of Gene Expression Systems in the *Drosophila* Head. Neuron. 2001;32(4):657–71.

80. Bahrami S, Drabløs F. Gene regulation in the immediate-early response process. Advances in Biological Regulation. 2016;62:37–49.

81. Hanson MA, Lemaitre B. New insights on Drosophila antimicrobial peptide function in host defense and beyond. Current Opinion in Immunology. 2020;62:22–30.

82. Tanji T, Hu X, Weber ANR, Ip YT. Toll and IMD Pathways Synergistically Activate an Innate Immune Response in *Drosophila melanogaster*. Molecular and Cellular Biology. 2007;27(12):4578.

83. Irving P, Troxler L, Heuer TS, Belvin M, Kopczynski C, Reichhart J-M, et al. A genome-wide analysis of immune responses in *Drosophila*. Proceedings of the National Academy of Sciences. 2001;98(26):15119.

84. Duneau D, Ferdy J-B, Revah J, Kondolf H, Ortiz GA, Lazzaro BP, et al. Stochastic variation in the initial phase of bacterial infection predicts the probability of survival in D. melanogaster. eLife. 2017;6:e28298.

85. Wolowczuk I, Verwaerde C, Viltart O, Delanoye A, Delacre M, Pot B, et al. Feeding our immune system: impact on metabolism. Clin Dev Immunol. 2008;2008:639803.

86. Chi W, Dao D, Lau TC, Henriksbo BD, Cavallari JF, Foley KP, et al. Bacterial Peptidoglycan Stimulates Adipocyte Lipolysis via NOD1. PLOS ONE. 2014;9(5):e97675.

87. Chambers MC, Song KH, Schneider DS. Listeria monocytogenes Infection Causes Metabolic Shifts in Drosophila melanogaster. PLOS ONE. 2012;7(12):e50679.

88. Krejčová G, Danielová A, Nedbalová P, Kazek M, Strych L, Chawla G, et al. Drosophila macrophages switch to aerobic glycolysis to mount effective antibacterial defense. eLife. 2019;8:e50414.

89. Clark Rebecca I, Tan Sharon WS, Péan Claire B, Roostalu U, Vivancos V, Bronda K, et al. MEF2 Is an In Vivo Immune-Metabolic Switch. Cell. 2013;155(2):435–47.

90. Eisen MB, Spellman PT, Brown PO, Botstein D. Cluster analysis and display of genome-wide expression patterns. Proceedings of the National Academy of Sciences of the United States of America. 1998;95(25):14863–8.

91. Im JH. Functional and Population Genetics of Drosophila Innate Immunity: Cornell University; 2018.

92. Lemaitre B, Hoffmann J. The Host Defense of Drosophila melanogaster. Annu Rev Immunol. 2007;25(1):697–743.

93. Konopka RJ, Benzer S. Clock Mutants of *Drosophila melanogaster*. Proceedings of the National Academy of Sciences. 1971;68(9):2112.

94. Myers MP, Wager-Smith K, Rothenfluh-Hilfiker A, Young MW. Light-induced degradation of TIMELESS and entrainment of the *Drosophila* circadian clock. Science. 1996;271(5256):1736–40.

95. McKean KA, Nunney L. Increased sexual activity reduces male immune function in Drosophila melanogaster. Proceedings of the National Academy of Sciences of the United States of America. 2001;98(14):7904–9.

96. Early AM, Shanmugarajah N, Buchon N, Clark AG. Drosophila Genotype Influences Commensal Bacterial Levels. PLOS ONE. 2017;12(1):e0170332.

97. Chambers MC, Jacobson E, Khalil S, Lazzaro BP. Thorax Injury Lowers Resistance to Infection in Drosophila melanogaster. Infection and Immunity. 2014;82(10):4380–9.

98. Sefer E, Kleyman M, Joseph Z-B. Tradeoffs between dense and replicate sampling strategies for high throughput time series experiments. Cell systems. 2016;3(1):35–42.

99. Robinson MD, Oshlack A. A scaling normalization method for differential expression analysis of RNA-seq data. Genome Biology. 2010;11(3):R25.

100. Mi H, Muruganujan A, Ebert D, Huang X, Thomas PD. PANTHER version 14: more genomes, a new PANTHER GO-slim and improvements in enrichment analysis tools. Nucleic Acids Research. 2018;47(D1):D419–D26.

101. Efron B, Tibshirani R. On testing the significance of sets of genes. Ann Appl Stat. 2007;1(1):107–29.

102. Subramanian A, Tamayo P, Mootha VK, Mukherjee S, Ebert BL, Gillette MA, et al. Gene set enrichment analysis: A knowledge-based approach for interpreting genome-wide expression profiles. Proceedings of the National Academy of Sciences. 2005;102(43):15545.

103. Mullighan CG, Su X, Zhang J, Radtke I, Phillips LAA, Miller CB, et al. Deletion of IKZF1 and prognosis in acute lymphoblastic leukemia. The New England journal of medicine. 2009;360(5):470–80.

104. Montero P, Vilar JA. TSclust: An R Package for Time Series Clustering. Journal of Statistical Software. 2014;62(1):1–43.

105. Galeano P, Peña D. Multivariate analysis in vector time series. Resenhas Do Instituto De Matemática E Estatística Da Universidade De São Paulo. 2000;4(4):383–404.

106. Mukhopadhyay ND, Chatterjee S. Causality and pathway search in microarray time series experiment. Bioinformatics. 2007;23(4):442–9.

107. Basu S, Shojaie A, Michailidis G. Network Granger Causality with Inherent Grouping Structure. Journal of Machine Learning Research. 2015;16:417–53.

